# Seasonality of pollinators in Mediterranean montane habitats: cool-blooded bees for early-blooming plants

**DOI:** 10.1101/2022.09.07.506992

**Authors:** Carlos M. Herrera, Alejandro Núñez, Luis O. Aguado, Conchita Alonso

## Abstract

Understanding the factors that drive community-wide assembly of plant-pollinator systems along environmental gradients has considerable evolutionary, ecological and applied significance. Variation in thermal environments combined with intrinsic differences among pollinators in thermal biology (tolerance limits, thermal optima, thermoregulatory ability) have been proposed as drivers of community-wide pollinator gradients, but this suggestion remains largely speculative. We test the hypothesis that seasonality in bee pollinator composition in montane habitats of southeastern Spain, which largely reflects the prevalence during the early flowering season of mining bees (*Andrena*), is a consequence of the latter’s thermal biology. Quantitative information on seasonality of *Andrena* bees in the whole plant community (275 plant species) was combined with field and laboratory data on key aspects of the thermal biology of 30 species of *Andrena* (endothermic ability, warming constant, relationships of body temperature with ambient and operative temperatures). *Andrena* bees were a conspicuous, albeit strongly seasonal component of the pollinator assemblage of the regional plant community, visiting flowers of 153 different plant species (57% of total). Proportion of *Andrena* relative to all bees reached a maximum among plant species which flowered in late winter and early spring, and declined precipitously from May onwards. *Andrena* were recorded only during the cooler segment of the annual range of air temperatures experienced at flowers by the whole bee assemblage. These patterns can be explained by features of *Andrena*’s thermal biology: null or negligible endothermy; ability to forage at much lower body temperature than endothermic bees (difference ~10°C); low upper tolerable limit of body temperature, beyond which thermal stress presumably precluded foraging at the warmest period of year; weak thermoregulatory capacity; and high warming constant enhancing ectothermic warming. Our results demonstrate the importance of lineage-specific pollinator traits as drivers of seasonality in community-wide pollinator composition; show that exploitation of cooler microclimates by bees does not require endothermy; falsify the frequent assumption that endothermy and thermoregulation apply to all bees; and suggest that medium- and large-sized ectothermic bees with low upper thermal limits and weak thermoregulatory ability can actually be more adversely affected by climate warming than large endothermic species.

## Introduction

Composition of pollinator assemblages at the plant community level often varies predictably along major ecological gradients such as plant cover, habitat type, elevation or time of year, such variation generally involving a turnover in the relative importance of different pollinators along the environmental axis considered (Cruden 1972, Kalin Arroyo et al. 1982, Shmida and Dukas 1990, McCabe et al. 2019, Herrera 2020, Dellinger et al. 2021). Some well-known examples of elevational or seasonal gradients in pollinator composition at the plant community level include the predominance of dipterans at high elevations or latitudes (Kalin Arroyo et al. 1982, Warren et al. 1988, Kearns 1992, Elberling and Olesen 1999, Hingston and McQuillan 2000, Koch et al. 2020); the shift from bee to vertebrate pollinators with increasing elevation in tropical mountains (Cruden 1972, Perillo et al. 2017, Dellinger et al. 2021); and the seasonal turnover of major orders and families of insect pollinators in Mediterranean plant communities (Shmida and Dukas 1990, Petanidou and Vokou 1993, Bosch et al. 1997). Since distinct groups of pollinators generally differ in crucial aspects of pollinating service which influence plant fitness and reproductive success (Schemske and Horvitz 1984, Herrera 1987, Valverde et al. 2019), understanding the causal mechanisms that drive the assembly of plant-pollinator systems along ecologically meaningful gradients has considerable evolutionary and ecological importance (Cruden 1972, Motten 1986, Dellinger et al. 2021, LaManna et al. 2021, Albor et al. 2022).

Differences between pollinator groups in thermal optima, thermal tolerance limits and/or thermoregulatory biology, acting in concert with spatial or temporal variation in abiotic factors (e.g., solar radiation, air temperature), have been traditionally the favored mechanistic explanation for elevational or seasonal gradients in pollinator composition (e.g., Warren et al. 1988, Shmida and Dukas 1990, Dellinger et al. 2021). Rather strikingly, however, no study has hitherto gone one step further beyond plausible explanations and actually undertaken comprehensive empirical work on the thermal biology of taxa involved in seasonal or elevational turnovers in pollinator composition. In addition to serving for a better understanding of the factors that drive the assembly of plant-pollinator systems, investigations on pollinator thermal biology have recently acquired special relevance because of the increasing concerns on the impact of climate change on pollinator populations, since the latter’s responses to accelerating changes in the thermal environment could possibly be contingent on their thermal biology (Herrera 1997, 2019, Ploquin et al. 2013, Scaven and Rafferty 2013, Marshall et al. 2018, Shrestha et al. 2018, Ghisbain et al. 2021). The central goal of this paper is to present an explicit test of the oft-mentioned but empirically untested hypothesis that the intrinsic thermal biology of pollinators can be a major driver of pollinator composition at the plant community level. We will focus on a community-wide seasonal pattern in bee pollinator composition that is apparently universal at mid and high latitudes in the Holarctic realm, namely the prevalence of mining bees (*Andrena*) during the early stages of the flowering season.

### *Seasonality of* Andrena *bees in the Holarctic realm*

Mining bees (Andrenidae) are a major family with >3,000 described species, ~1600 of which are in the predominantly Holarctic genus *Andrena*, the second largest genus of bees (Bossert et al. 2022, Pisanty et al. 2022). *Andrena* bees are pollinators of many wild plants in a broad variety of boreal and temperate habitats of Eurasia and North America, particularly of species flowering in winter and early spring (Schemske et al. 1978, Motten et al. 1981, Anderson and Beare 1983, Motten 1986, Kato 2000, Ostaff et al. 2015, Turley et al. 2022, Wood et al. 2022). The association between early blooming and *Andrena* pollination stems from the prevailing phenology of these bees, which are among the earliest-flying ones in temperate and boreal habitats, can nest when it is often cold and rainy, and are able to withstand thermally unfavorable climates, as illustrated by their abundance and diversity in arctic and subarctic habitats (Sakagami and Matsumura 1967, Armbruster and Guinn 1989, Batra 1990, Kato 2000, Hicks and Sheffield 2021).

Studies on the pollination ecology of early-blooming plants have mainly focused on the reproductive features of plants (phenology, floral biology, pollen limitation; Schemske et al. 1978, Motten et al. 1981), yet the thermal characteristics of their associated *Andrena* pollinators have been addressed on few occasions. One of these studies showed that pollination of the early-flowering daffodil *Narcissus longispathus* was facilitated by the thermal biology of *Andrena bicolor*, its main pollinator, which flies at low body temperature and has a low upper thermal tolerance limit in comparison to other bees (Herrera 1995). It remains unknown, however, whether these distinctive thermal features apply to the genus *Andrena* as a whole, as knowledge on bee thermal biology largely refers to a few lineages of endothermic bees in the family Apidae (Inouye 1975, Chappell 1982, May and Casey 1983, Stone and Willmer 1989, Stone 1993, Heinrich 1993, Roberts et al. 1998). The thermal biology of *Andrena* bees remains essentially unexplored (Danforth et al. 2019; but see Herrera 1995, Schmaranzer et al. 1997, Bishop and Armbruster 1999), which represents a remarkable gap in our knowledge given their extraordinary biological diversity, broad geographical distribution, high seasonal abundance, and importance as pollinators of many cultivated and wild plants. In particular, it is unknown whether the characteristic early flying period of *Andrena* bees, which enhances their role as pollinators of early-blooming plants can be explained by their thermal biology.

We present in this paper a quantitative analysis of the seasonal occurrence and thermal environment of *Andrena* pollinators in a montane area of southeastern Spain, along with observational and experimental evidence on the thermal biology of *Andrena* bees obtained in the field and the laboratory. The following specific questions will be addressed: (1) How does the proportional abundance of *Andrena* pollinators relative to other bees varies seasonally ?; (2) Does the thermal environment experienced by foraging *Andrena* bees differ from that of other bees ?; (3) To what degree does body temperature of *Andrena* bees in the field depend on ambient temperature and operative body temperature (*sensu* Bakken 1992, see definition later), and which are the shapes of the relationships ?; and (4) Are *Andrena* individuals able to warm by themselves through endothermy in absence of external heat sources ? Questions (1) and (2) will be addressed for the whole regional community of insect-pollinated plants and their bee pollinators, while questions (3) and (4) will be answered for a taxonomically diverse sample of species comprising a substantial fraction of the *Andrena* occurring in the region studied. Responses to these specific questions will allow to evaluate the hypothesis that *Andrena* seasonality is a consequence of its thermal biology. More generally, our results will bear on the role of pollinators’ thermal biology as a driver of seasonal community-wide variations in pollinator composition and on the expected impact of climate warming on bee pollinator assemblages.

## Materials and methods

### Study area and sampling periods

Data on bee pollinator composition and air temperature at the foraging sites of different bee species were collected during January-December 1997-2022 in a relatively small area of the Sierra de Cazorla, Jaén Province, southeastern Spain (see Herrera 2021: Fig. 2, for map of sampling sites). This region is characterized by their well-preserved montane habitats and outstanding biological diversity (Médail and Diadema 2009, Gómez Mercado 2011). All *Andrena* bees used in laboratory experiments, and most of those whose thoracic temperatures were measured in the field, were also caught there during February-May 1994-2022. This seasonal sampling schedule closely matched the main activity period of *Andrena* species in the region (see Results). To increase sample size for some species that are infrequent in Sierra de Cazorla, or to augment the number of subgenera and hence the taxonomic breadth represented in the sample, additional field measurements of *Andrena* thoracic temperature were gathered in February-April 2022 at two locations in Sierra Morena (Córdoba province) and the lowlands of the Guadalquivir River valley (Sevilla province), 160 km and 300 km away, respectively, from the Cazorla main study area.

### Bee pollinators and their thermal microenvironment

Pollinator composition was quantitatively assessed for 275 plant species in 179 genera from 47 families (see Appendix S1: Table S1 for species list). This sample is a superset of the 221 species studied by Herrera (2021), and includes virtually all entomophilous plants occurring in the area of the Sierra de Cazorla sampled for bees. Mean pollinator sampling date for each plant species roughly matched its peak flowering date, hence the seasonal distribution of sampling times closely matched the pattern of flowering times in the region (Appendix S1: Table S1). About 75% of plant species in the sample (*N* = 208) were sampled for pollinators only one year, and the rest were sampled on ≥ 2 years. Pollinator sampling was conducted on a single site in the vast majority of the species considered here (*N* = 258), while 17 species were sampled on two or more sites. Pollinators of some species considered here vary among sites or years, but such intraspecific variation is quantitatively minor in comparison to the broad interspecific range occurring in the large species sample considered here, as previously shown by Herrera (2020: Table 2, Fig. 2) by means of formal variance partitions. Pollinator data for the same plant species obtained in more than one site or year were thus combined into a single sample for the analyses. The quality of pollinator composition data, as assessed with Good-Turing’s “sample coverage” parameter (= estimate of the proportion of the total population of pollinators that is represented by the species occurring in the sample; Hsieh et al. 2016), was very high (>85%) in the vast majority of plant species (Herrera 2021).

The proportions of *Andrena* and non-*Andrena* bees were obtained for each plant species using quantitative data on pollinator composition obtained using the field methods described in detail by Herrera (2019, 2020). The elemental sampling unit was the “pollinator census”, consisting of a 3min watch of a flowering patch whose total number of open flowers was also counted. All pollinators visiting some flower in the focal patch during the 3min period were identified. For all plant species combined, a total of 33,446 censuses were conducted on 696 different dates, yielding a total of 1,431 *Andrena* and 11,386 non-*Andrena* bee individuals (see Appendix S1: Table S1 for detailed data on individual plant species).

Air temperature at bee foraging sites was measured over January-August for a total of 1,327 bees from 58 species (19 *Andrena*, 39 non-*Andrena*) and 19 genera while they were visiting flowers of 18 different plant species in fine weather (see Appendix S1: Table S2 for bee species and number of measurements). Measurements were taken in a subset of the localities where pollinator censuses were carried out, and were representative of all major habitat types occurring in the region. Foraging bees were hand netted, identified to species (or collected to be identified later), and air temperature (*T*_a_ hereafter) at the site where the bee had been caught was measured shortly after the bee’s capture using a digital thermometer and a fast-response 0.22 mm-diameter thermocouple. These data will be used here to compare the thermal profiles of the foraging sites of *Andrena* and non-*Andrena* bees on an annual basis (*N* = 328 and 999 measurements, respectively).

### *Thermal biology of* Andrena

Data were obtained in the laboratory and/or the field on the thermal biology of 30 species of *Andrena* belonging to 16 different subgenera (Table 1). They represent 40% of the ~75 species of *Andrena* recorded in the study area during the study period (C. M. Herrera, A. Núñez and C. Alonso, *personal observations*), and encompass the whole range of body sizes occurring there excepting those in the subgenus *Micrandrena*, whose minute body sizes precluded the application of the methods used in this investigation.

**Table 1.** The 30 species of *Andrena* considered in this study, listed in alphabetical order of subgenera.

| Species | Subgenus | Part of the study |  |
| --- | --- | --- | --- |
|  |  | Field<br>measurements | Laboratory<br>experiments |
| *Andrena fulva* (Müller, 1766) | *Andrena* | X | X |
| *Andrena lagopus* Latreille, 1809 | *Biareolina* | X |  |
| *Andrena humilis* Imhoff, 1832 | *Chlorandrena* |  | X |
| *Andrena livens* Pérez, 1895 | *Chlorandrena* | X | X |
| *Andrena rhenana* Stöckhert, 1930 | *Chlorandrena* | X |  |
| *Andrena rhyssonota* Pérez, 1895 | *Chlorandrena* | X | X |
| *Andrena bicolor* Fabricius, 1775 | *Euandrena* | X | X |
| *Andrena vulpecula* Kriechbaumer, 1873 | *Euandrena* |  | X |
| *Andrena labialis* (Kirby, 1802) | *Holandrena* | X | X |
| *Andrena trimmerana* (Kirby, 1802) | *Hoplandrena* | X | X |
| *Andrena baetica* Wood, 2020 | *Lepidandrena* | X |  |
| *Andrena sardoa* Lepeletier, 1841 | *Lepidandrena* | X | X |
| *Andrena albopunctata* (Rossi, 1792) | *Melandrena* |  | X |
| *Andrena assimilis* Radoszkowski, 1876 | *Melandrena* | X | X |
| *Andrena hispania* Warncke, 1967 | *Melandrena* |  | X |
| *Andrena nigroaenea* (Kirby, 1802) | *Melandrena* | X | X |
| *Andrena thoracica* (Fabricius, 1775) | *Melandrena* |  | X |
| *Andrena schencki* Morawitz, 1866 | Opandrena |  | X |
| *Andrena pilipes* Fabricius, 1781 | Plastandrena |  | X |
| *Andrena tibialis* (Kirby, 1802) | Plastandrena |  | X |
| *Andrena impressa* Warncke, 1967 | Ptilandrena |  | X |
| *Andrena antigana* Pérez, 1895 | Simandrena | X |  |
| *Andrena combinata* (Christ, 1791) | Simandrena |  | X |
| *Andrena congruens* Schmiedeknecht, 1884 | Simandrena |  | X |
| *Andrena lepida* Schenck, 1861 | Simandrena |  | X |
| *Andrena ovatula* (Kirby, 1802) | Taeniandrena |  | X |
| *Andrena haemorrhoa* (Fabricius, 1781) | Trachandrena |  | X |
| *Andrena ferrugineicrus* Dours, 1872 | Truncandrena |  | X |
| *Andrena truncatilabris* Morawitz, 1877 | Truncandrena | X |  |
| *Andrena flavipes* Panzer, 1799 | Zonandrena | X | X |

#### Laboratory experiments

The ability of live *Andrena* bees to warm-up autonomously by endothermy, and the intrinsic warming constants of the same individuals, were assessed experimentally. Bee specimens for experiments were netted in the field while they were foraging at flowers, placed into sealed microcentrifuge tubes kept in the dark in an ice bath, and quickly brought to the laboratory. All measurements were made within 4 h of capture. The junction of a 0.22mm Type T thermocouple was implanted in the thorax to a depth of 1 mm, and held in place using a small amount of a wax-resin mixture. The bee was briefly cooled in a refrigerator until its thoracic temperature (*T*_th_ hereafter) was about 8°C, and then placed on a piece of styrofoam in a small room with still air and without any source of radiation (room temperature 18-21°C). *T*_th_ and air temperature 10 cm away from the bee were continously recorded every 10 s using a data logger. The bee was allowed to warm up autonomously until *T*_th_ stabilized. During this period its abdomen was gently pinched with forceps to test for flight ability. After *T*_th_ stabilized for at least 30 s, the bee was killed, weighed to the nearest 0.1 mg and briefly cooled in a refrigerator. The dead bee was then quickly placed again on the same styrofoam piece and allowed to warm up spontaneously until *T*_th_ stabilization. As with the live bee, *T*_th_ and air temperature 10 cm away from the bee were recorded every 10 s with a data logger. Representative examples of experimental warming curves for live and dead experimental specimens of eight species differing in warming ability are shown in Fig. 1. This experimental protocol was applied to 125 individuals from 25 *Andrena* species, and the data obtained will be used to obtain quantitative estimates of their warming ability and warming constants (see *Data analyses* below).

**Fig. 1.**
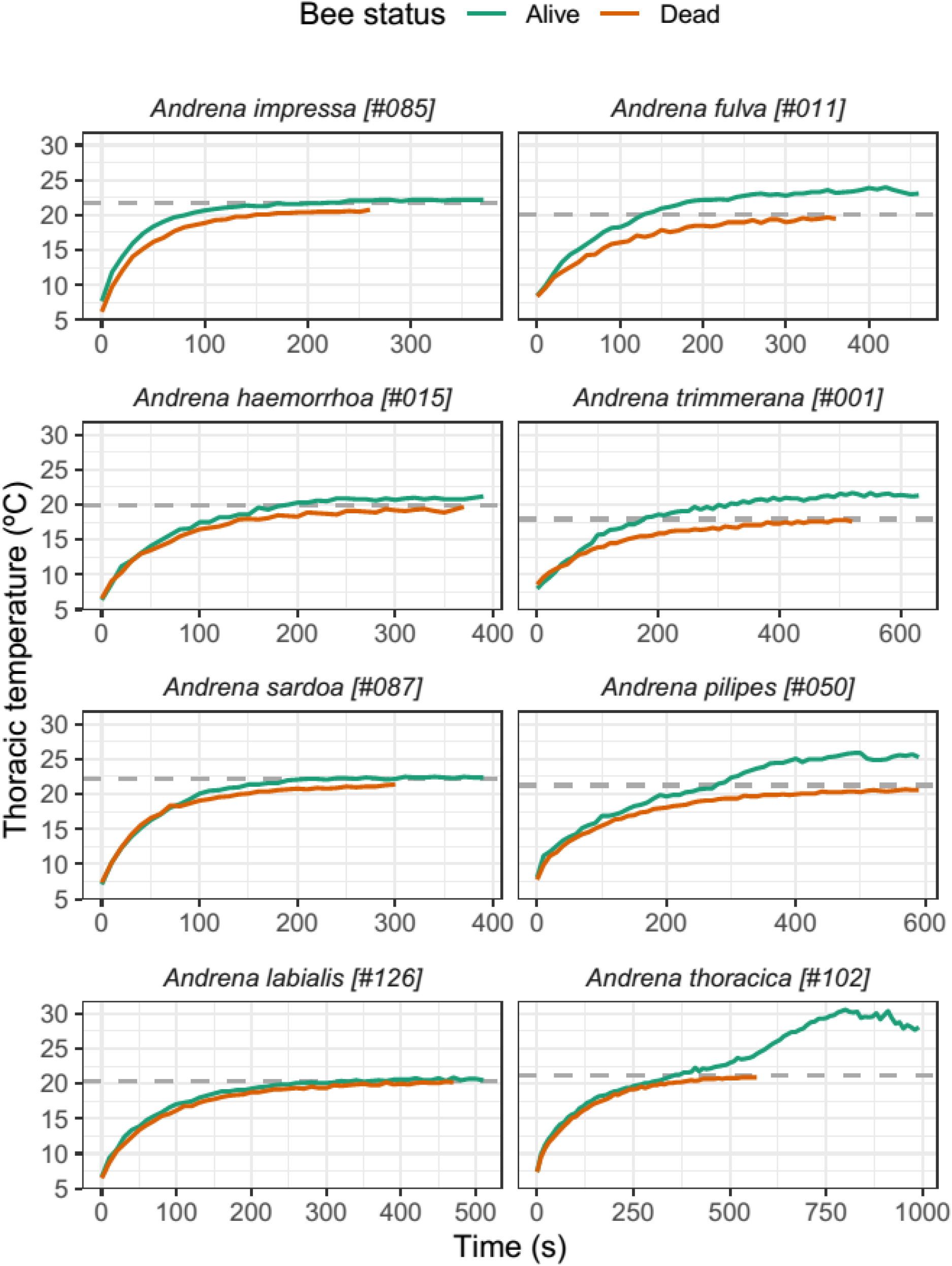
Experimental warming curves for eight individuals of different *Andrena* species, chosen to illustrate the range of variation in autonomous endothermy represented in the sample. Individuals in the left and right columns exhibited null and weak endothermic ability, respectively. Horizontal scales vary among graphs. Horizontal dashed lines denote the mean ambient temperature during each experimental run. Numbers in brackets following species names are specimen identifiers.

#### Field measurements

Free-ranging individuals of *Andrena* species were hand netted in fine weather while foraging at flowers of different species. Efforts were made to sample bees over as broad ranges as possible of habitat types and ambient temperature. For each bee, *T*_th_ was measured within 10 s of netting to the nearest 0.1°C using a fast-response (time constant 0.025 s), 0.33 mm-diameter needle microprobe with sharpened tip (Type MT-29/1; Physitemp Instruments, Clifton, New Jersey, USA). Readings were obtained by inserting the probe 1 mm into the bee’s thorax dorsally while it was restrained in the net. Air temperature (*T*_a_) at the same spot where the bee had been caught was measured within 2 min of the bee’s capture using a digital thermometer and a fast-response 0.22 mm-diameter thermocouple. A total of 497 paired *T*_th_ - *T*_a_ measurements on foraging individuals of 15 *Andrena* species were obtained in the field on 42 different sampling dates.

A subset of the individuals with paired *T*_th_ and *T*_a_ field data (*N* = 186 individuals from seven species) were killed immediately after thoracic temperature measurement using ethyl acetate, and their “perception” of the natural thermal environment was assessed by measuring their operative temperatures (*T*_e_). *T*_e_ represents the equilibrium temperature that an organism would reach under stable conditions in the absence of metabolic input, and it simultaneously reflects air temperature and a temperature increment or decrement subsuming radiative and convective factors (Bakken 1989, 1992, Bishop and Armbruster 1999). The needle thermocouple was inserted into the thorax of the freshly-killed bee, which was then placed lying flat on the surface of the same flower on which it had been previously captured. *T*_th_ was subsequently recorded with a digital thermometer until variation became negligible (<0.5°C in 30 s), and this final value was used as an estimate of *T*_e_. All individuals used for *T*_e_ estimation were weighed to the nearest 0.1 mg within a few hours of measurements.

### Data analyses

All statistical analyses reported in this paper were carried out using the R environment (R Core Team, 2020). The proportional importance of *Andrena* bees relative to all bees combined (Andrenidae, Apidae, Colletidae, Halictidae and Megachilidae) was estimated for each plant species by dividing the total number of *Andrena* individuals recorded in censuses by the total number of individuals for all bees combined (see Appendix S1: Table S1 for raw data and estimated proportions). Nine plant species without any bee pollinator were excluded from all analyses. The shape of the seasonal trend in the relative importance of *Andrena* as pollinators in the study region was assessed by regressing the proportion of *Andrena* for every plant species (*N* = 266) against the mean census date for the species (expressed as days from 1 January), using the generalized additive model smoother implemented in function gam of the mgcv package for R (Wood 2017).

Since many *Andrena* species are oligolectic on one or a few phylogenetically related plant species (Larkin et al. 2008, Wood and Roberts 2018) and the flowering period of plant species is known to have a strong phylogenetic signal (Kochmer and Handel 1986, Davies et al. 2013), then *Andrena* seasonality could actually depend more on their association with certain groups of plants which happen to flower during a particular period of year rather than on the particular period of the year itself. To analytically address this possibility, statistical significance of the seasonal trend in frequency of *Andrena* bees was evaluated by fitting two separate generalized linear mixed models with *Andrena* proportion for each plant species as the response variable, mean census date for the species as the single fixed effect, and either plant family or plant genus as random effects. This approach allowed to dissect the effects of time of year and plant taxonomical composition on *Andrena* frequency. Computations were performed with the glmer function of the lme4 package (Bates et al. 2015), the response variable was modelled as a binomial process, and dates (expressed as days from 1 January) were scaled to mean zero and standard deviation unit, so that fixed-effect parameter estimates represented standardized effects. Confidence intervals for model parameter and variance estimates were obtained using function confint. merMod.

Laboratory runs on *Andrena* bees yielded estimates of two thermal parameters for each tested individual: the maximum thermal excess relative to ambient temperature reached autonomously by the living bee and maintained during at least 30 s (*T*_exc_ = *T*_th_ – *T*_a_); and the constant *K* describing heat transfer rate obtained by fitting the passive warming curve of the dead bee to the equation *dT/dt* = *K*(*T-T*_a_ (Newton’s law of cooling; Willmer and Unwin 1981; Casey 1988), in which *T* is the temperature of the object, *t* is time, *T*_a_ is ambient temperature, and *K* is the cooling/warming constant (“warming constant” will be used hereafter for brevity). The nonlinear least-squares method implemented in the function nls of the stats package was used to estimate *K* from empirical experimental data. *T*_exc_ and *K* values will be used to describe the warming ability and inherent heat transfer rates of the species tested, respectively.

The functional relationships linking either thoracic (*T*_th_) or operative (*T*_e_) temperatures with ambient temperature (*T*_a_) under natural field conditions were assessed by regressing *T*_th_ or *T*_e_ against *T*_a_ separately for each species. As we were interested in assessing the shape of the functional relationships between temperatures and their possible departures from linearity, nonparametric regressions based on the generalized additive model smoother in function gam of the mgcv package were used (Wood 2017). This method, in contrast to ordinary parametric linear or quadratic regressions, does not make *a priori* assumptions on the shape on the statistical relationship between predictor and response variables.

Results obtained here for two key thermal features of *Andrena* bees (warming constant *K* estimated in the laboratory, and thoracic temperatures of foraging individuals measured in the field) were compared with those obtained in the study region for a sample of 20 bee species from other families and genera using identical field and laboratory procedures (Herrera 1997, and C. M. Herrera *unpublished observations*). A list of the species used in the comparisons along with the raw data and their sources are presented in Appendix S2: Table S1. The statistical relationships between *K* and body mass (both log-transformed), and between mean thoracic temperature and mean air temperature, for *Andrena* and non*Andrena* bees were compared by fitting two simple linear models to the data which included bee group, either body mass or air temperature, and the corresponding interaction term, as predictors. Fits and parameters estimates were obtained with the function lm of the stats package.

## Results

### *Distribution of* Andrena *bees over the seasonal and air temperature gradients*

*Andrena* bees were important pollinators in the regional assemblage of entomophilous plants. They were recorded visiting flowers of 153 different plant species, or 57.5% of species with flowers visited by bees (*N* = 266; Appendix S1: Table S1). Their proportional importance relative to all bees varied widely among plant species, and exhibited a distinct seasonal trend. The proportion of *Andrena* individuals in pollinator censuses tended to be highest among plant species flowering during February-May (Fig. 2). Individuals of *Andrena* contributed >50% of all bees in 29 of the 137 species flowering during that period (Fig. 2), and were the exclusive bee pollinators of some very early-blooming herbs such as six species of Brassicaceae in the genera *Alyssum*, *Erophila*, *Iberis*, *Jonopsidium* and *Lepidium* (Appendix S1: Table S1). The proportion of *Andrena* in the bee pollinator assemblages of individual plant species tended to decline precipitously from May onwards, remaining unrecorded or having negligible importance for plant species which flowered from late June through October (Fig. 2). The effect of time of year on the frequency of *Andrena* in the pollinator assemblages of individual plant species remained statistically significant after controlling for the variance among plant genera and families in flowering phenology (Table 2), thus indicating that *Andrena* seasonality was not an spurious consequence of their association with early-flowering taxa.

**Fig. 2.**
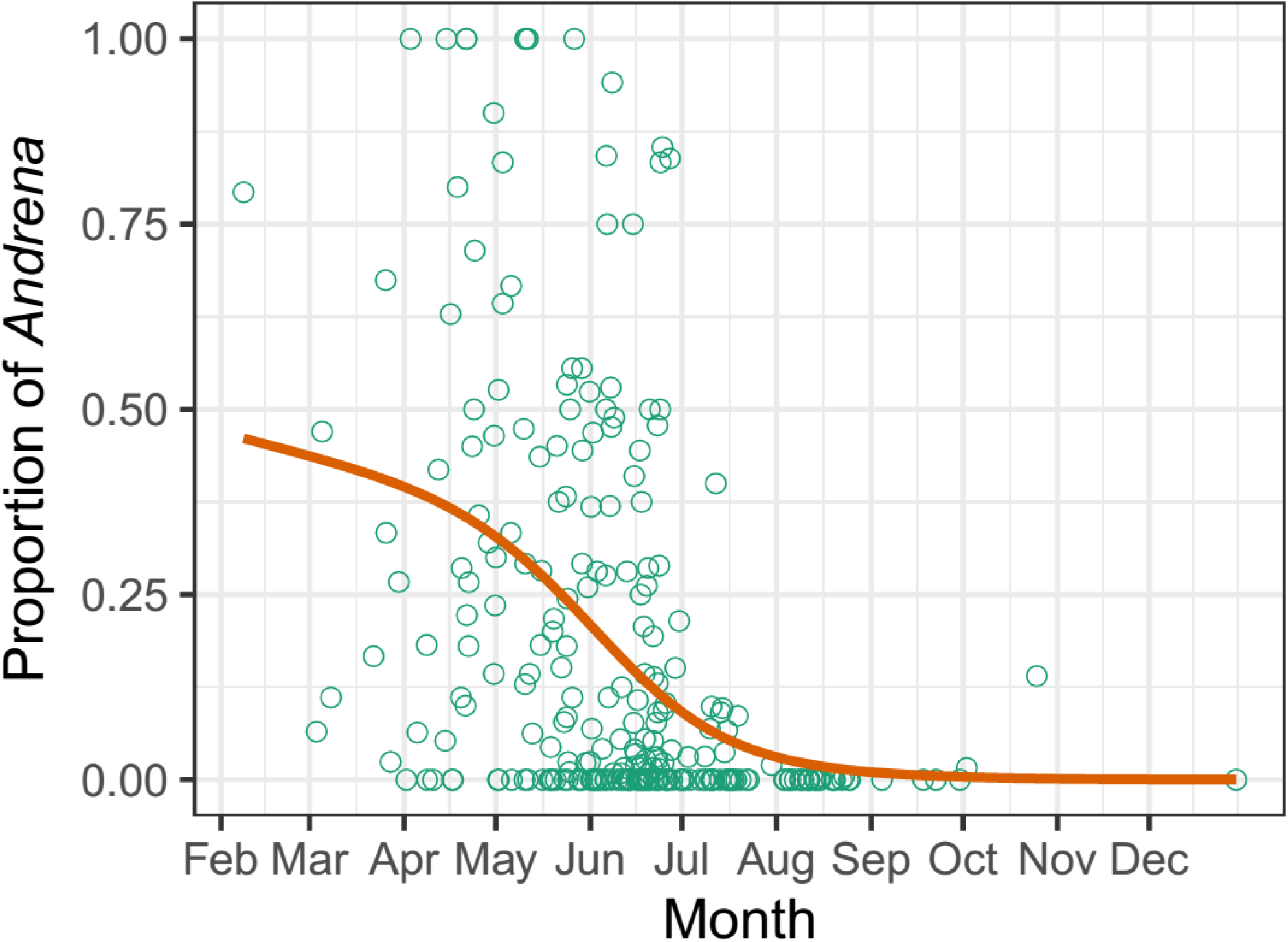
Seasonal variation in the proportion of *Andrena* relative to total bees in the pollinator assemblages of 266 species of bee-pollinated plants from montane habitats in the Sierra de Cazorla study area. Each symbol represents a different plant species. The horizontal axis is the mean census date for the species, and the vertical axis the proportion of bee individuals in pollinator censuses contributed by species of *Andrena*. The red line is a nonparametric regression fitted to the data using a binomial smoothing spline. See Appendix S1: Table S1 for a list of plant species, sampling effort, and the raw data plotted here.

**Table 2.**
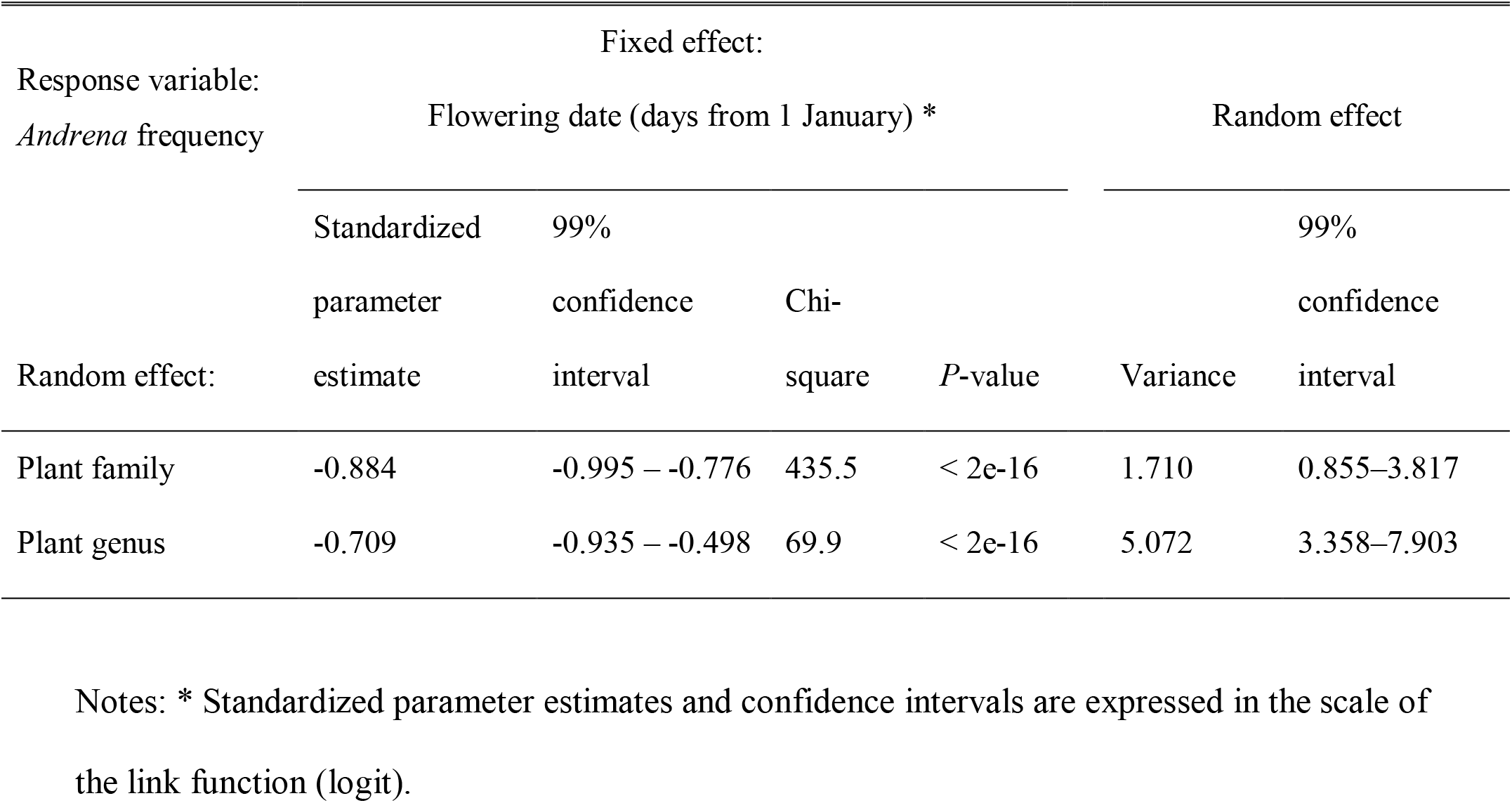
Summary of results of generalized linear mixed models testing for the effect of flowering date on the frequency of *Andrena* relative to all bees while controlling for differences in blooming time among plant genera and families (*N* = 266 plant species).

On an annual basis, *Andrena* and non-*Andrena* bees tended to be associated with contrasting thermal microenvironments while foraging, as revealed by the contrasting frequency distributions of air temperatures at capture points (Fig. 3). Both the central trend and the spread of the distribution were lower for *Andrena* (interquartile range = 16-22°C) than non-*Andrena* bees (interquartile range = 20-31°C). In addition, the upper limit of air temperatures at foraging sites was considerably lower for *Andrena*, as denoted by the observation that only 13% of all *Andrena* were found foraging at sites with air temperatures above the median of the combined distribution for all bees (23.3°C), while the corresponding figure for non-*Andrena* bees was 62% (Fig. 3).

**Fig. 3.**
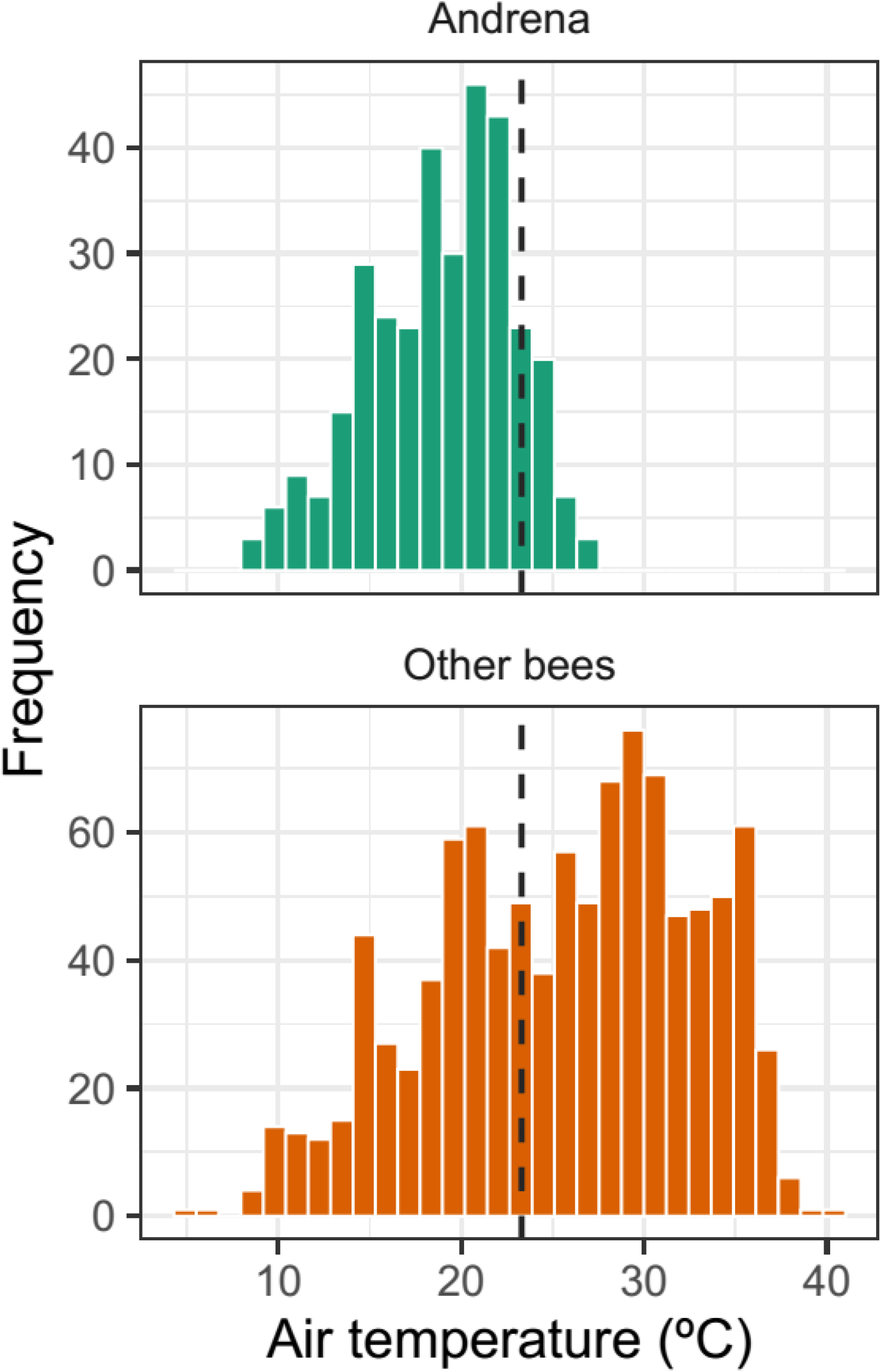
Frequency distributions of air temperature at foraging sites of *Andrena* and non-*Andrena* bees sampled throughout the year in the Sierra de Cazorla study area. The median value for the combined distribution of the two groups (23.3°C) is shown for reference (vertical dashed line).

### Endothermic ability and warming constant

Laboratory experiments revealed from null to weak endothermic ability in the sample of 25 *Andrena* species tested (Table 3). The autonomous increase in thoracic temperature above the ambient ranged between <1°C (e.g., *A. impressa, A. sardoa, A. livens*) and ~3°C (e.g., *A. nigroaenea*, *A. pilipes*, *A. thoracica*). There was a close positive linear relationship across species between mean *T*_exc_ and mean body mass (*r* = 0.791, *N* = 25, *P* = 2.5e-06; Fig. 4A). The endothermic ability of all small species (<50 mg body mass) was consistently null or negligible (mean *T*_exc_ 0-0.5°C), while only some of the largest-sized species had mean *T*_exc_ exceeding 3.0°C (Table 3; see also Fig. 1 for some individual examples). The warming constant *K* ranged between 0.0071-0.0141 s^−1^ in the set of species studied (Table 3), and this parameter was closely, inversely correlated with body mass across species (*r* = −0.845, *N* = 25, *P* = 1.1e07; Fig. 4B).

**Table 3.**
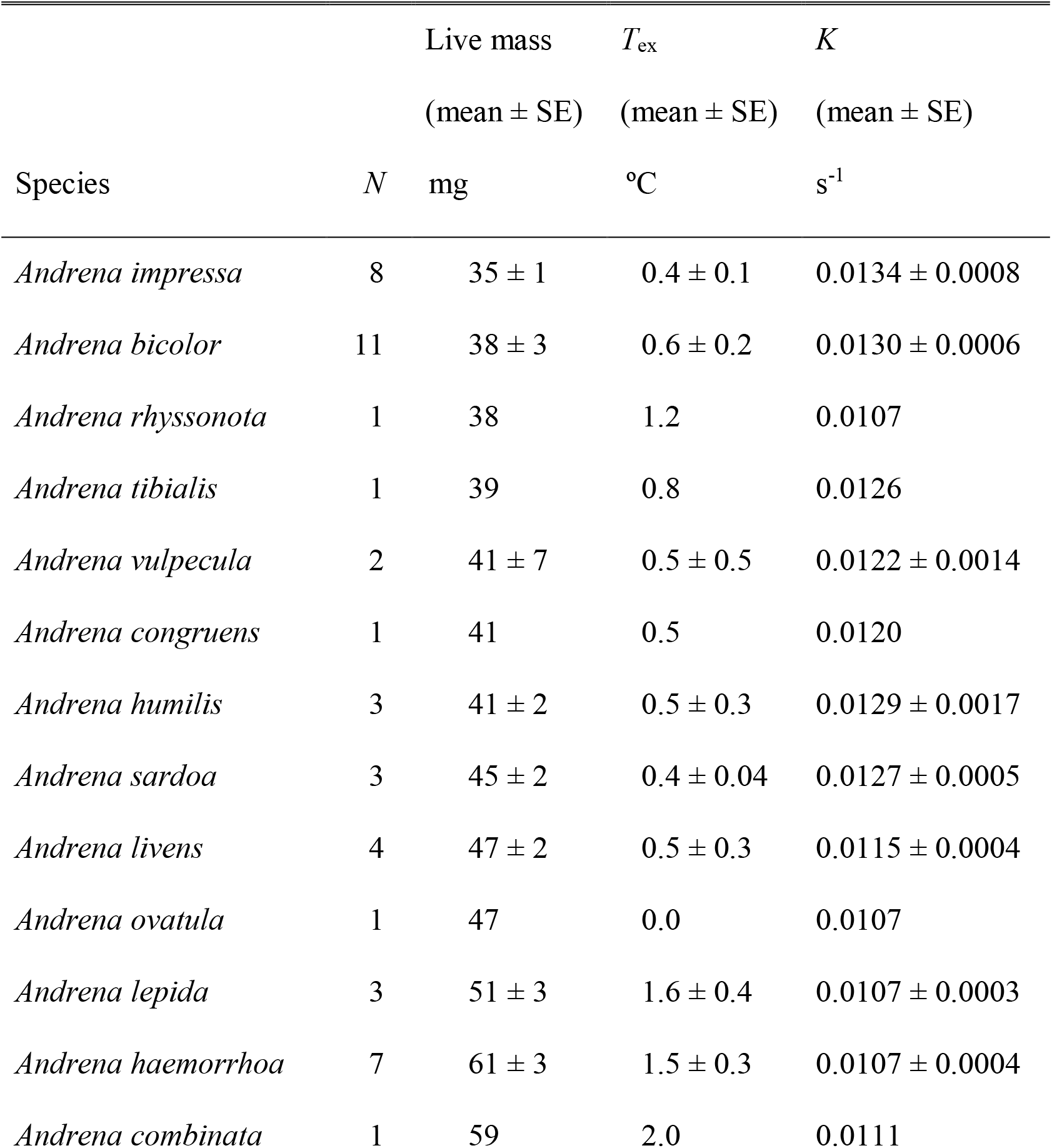
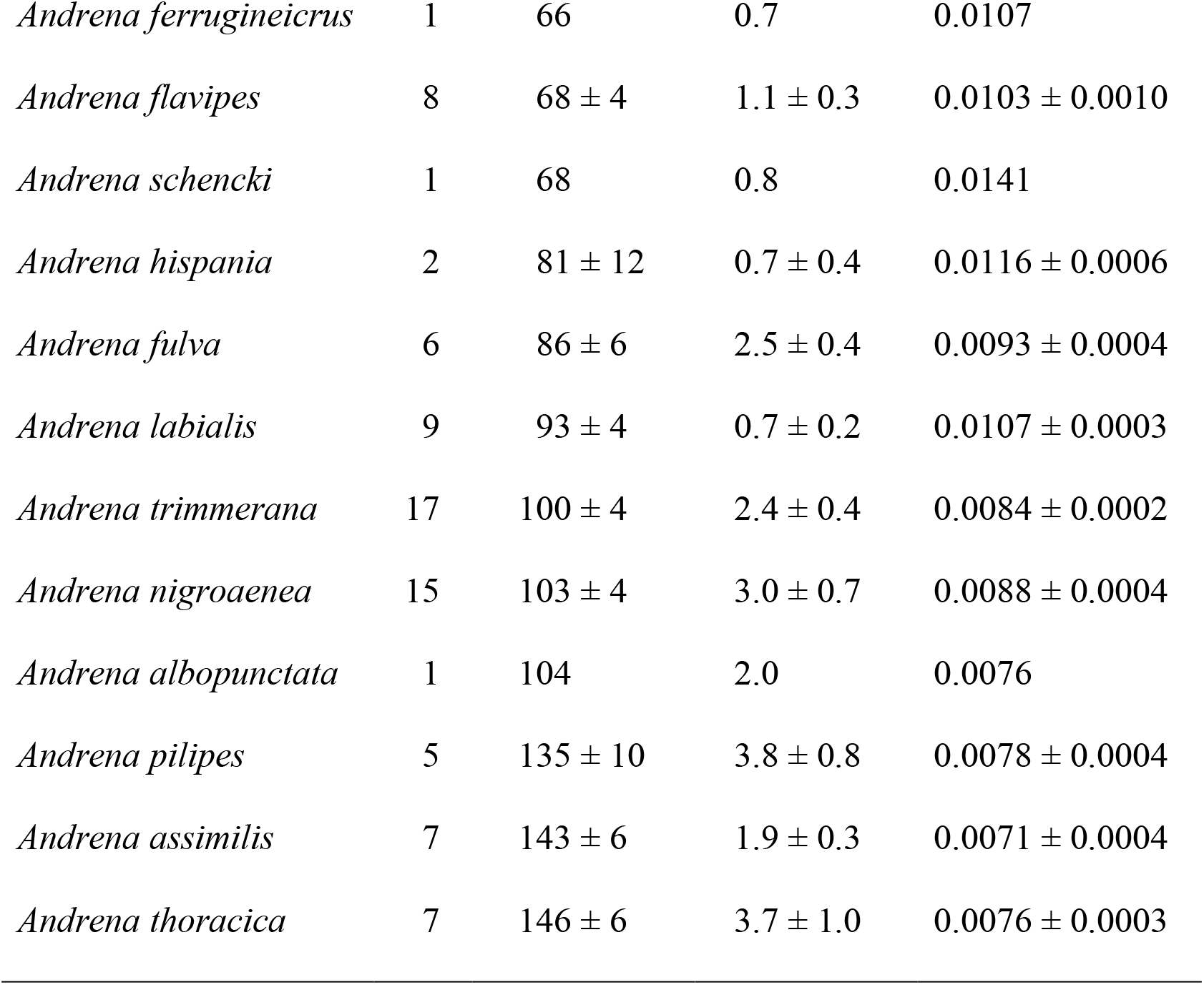
Mean body mass and thermal parameters for 25 species of *Andrena* tested experimentally in the laboratory. Species are listed in increasing order of body mass. *N* = number of individual bees tested; *T*_ex_ = maximum difference between thoracic and ambient temperature reached by live bees by spontaneous endothermy and maintained for at least 30 s; *K* = warming constant for dead bees, estimated by fitting to the equation for Newton’s law of cooling the observed temperature sequences of bees as they equilibrated thermally with the ambient.

**Fig. 4.**
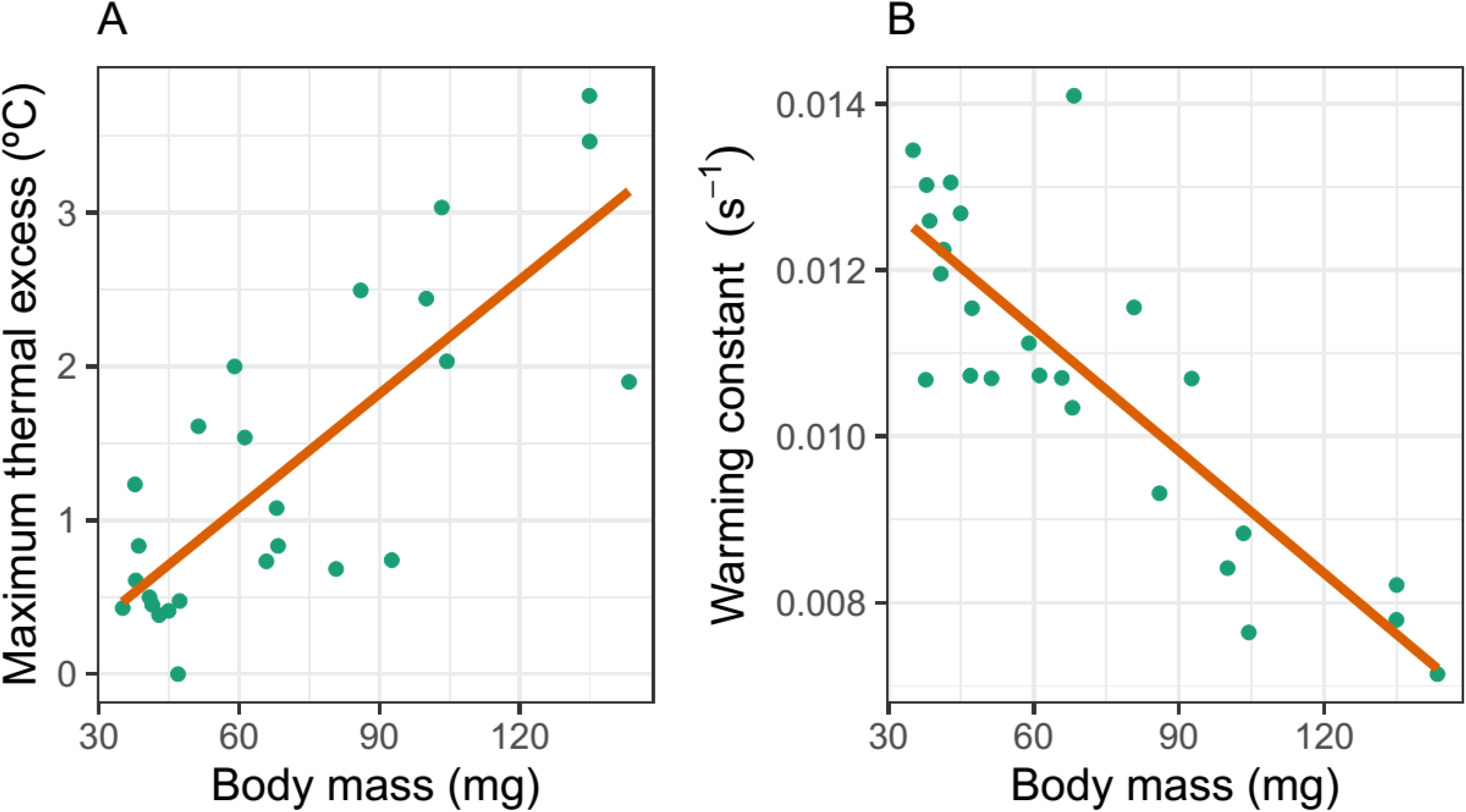
Relationships across *Andrena* species (*N* = 25) of endothermic ability (A) and warming constant (B) with mean body mass, as determined experimentally in the laboratory. Lines are nonparametric regressions fitted to the data using a generalized additive model and cubic splines, which implies that linearity of relationships was not a condition imposed *a priori* on the data by the fitting method.

### Thoracic temperature in relation to ambient temperature

Mean *T*_th_ for the 15 *Andrena* species sampled in the field are summarized in Fig. 5. There was little interspecific variation in mean *T*_th_ values, which ranged between 25.6°C (*A. livens*) and 31.3°C (*A. labialis*). Interspecific variation in mean *T*_th_ was not significantly correlated with variation in mean body mass (*r* = 0.112, df = 13, *P* = 0.69).

**Fig. 5.**
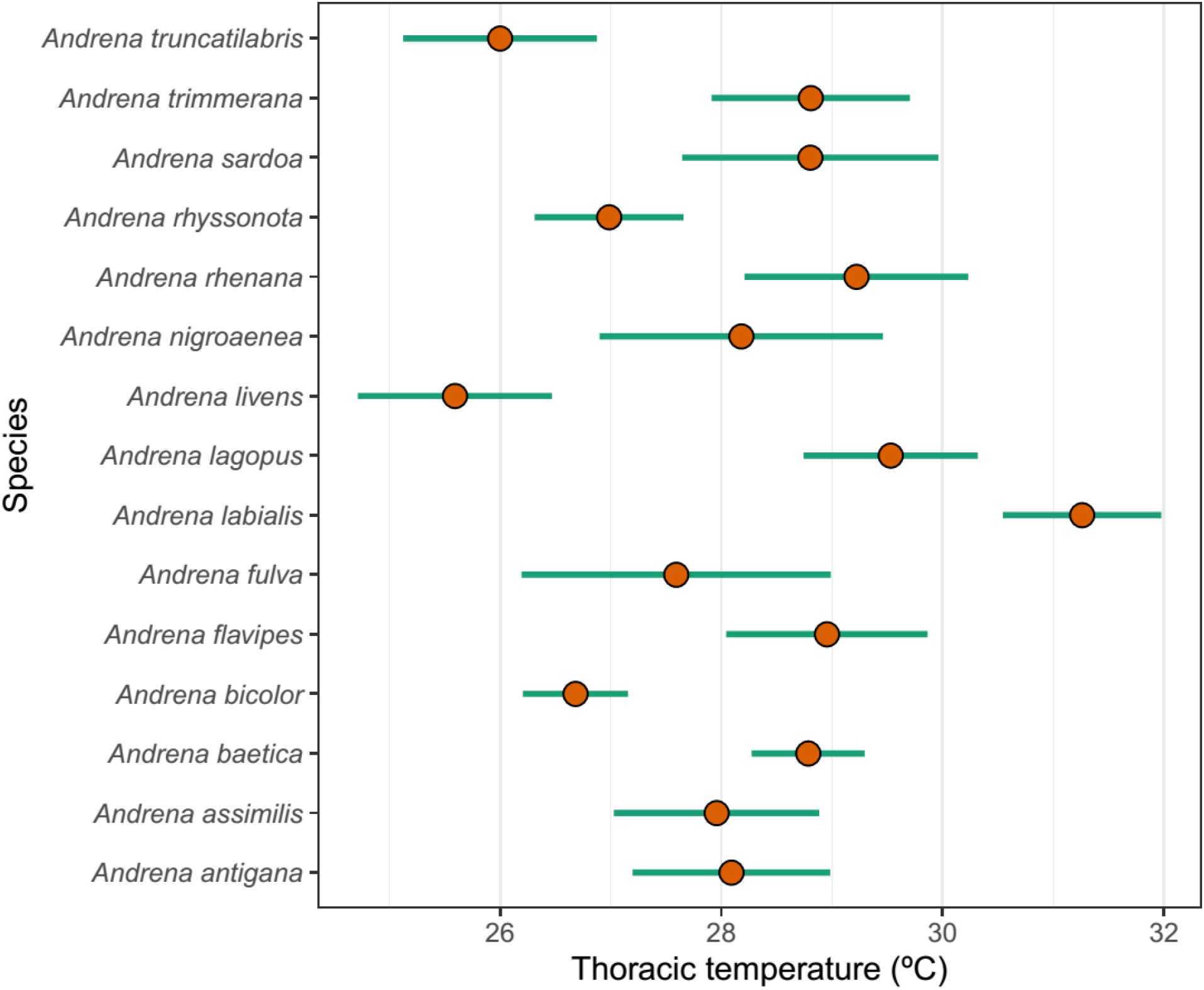
Mean thoracic temperature of foraging individuals of 15 species of *Andrena* measured in the field. Dots are means and segments extend over ± 2 standard errors.

All individual *Andrena* bees sampled in the field while foraging at flowers were substantially warmer than the air at the capture site, and *T*_th_ was significantly, positively related to *T*_a_ in all species (Fig. 6). The shape of the relationship between *T*_th_ and *T*_a_ was essentially linear in most species despite the fitting method used (generalized additive model smoother) not imposing linearity *a priori* on the data. Only in a few instances (*A. lagopus*, *A. livens*, *A. sardoa*) there was some evidence of weakly nonlinear, convex relationships (Fig. 6). In general, regressions tended to run parallel to the line *y* = *x*, thus denoting that variation in *T*_th_ closely followed variation in ambient temperature.

**Fig. 6.**
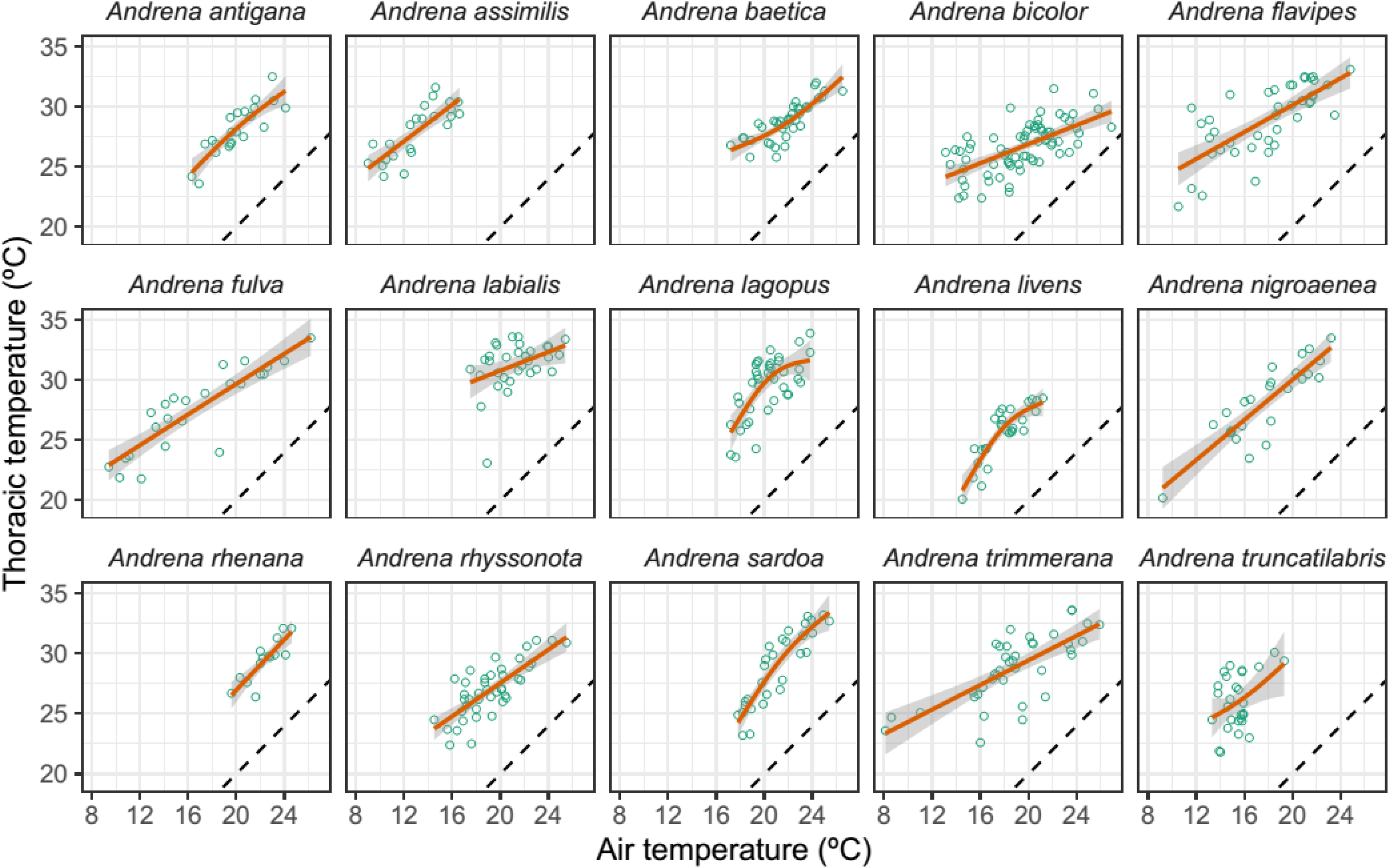
Relationship between bee thoracic temperature and air temperature at capture spot for 15 *Andrena* species sampled in the field. Each symbol corresponds to an individual bee, red solid lines are nonparametric regressions fitted using smoothing splines, and the shaded bands are 95% confidence envelopes around the fit. The dashed line depicts the *y* = *x* line.

### Thoracic temperature in relation to operative temperature

Thoracic and operative temperatures (*T*_th_ and *T*_e_, respectively) were measured concurrently on 186 individuals of seven species (*A. assimilis, A. flavipes, A. fulva, A. labialis, A. nigroaenea, A. sardoa, A. trimmerana*) that roughly encompassed the whole range of body mass and endothermic ability represented in the species sample considered in this study (Table 3). Despite differences in body size and endothermic ability, the shape of nonparametric fits of *T*_th_ against *T*_e_ were conspicuously similar in all species. Thoracic temperature was positively, nonlinearly related to operative temperature (Fig. 7). The fitted curves for all species consistently intersected the *y* = *x* line at *T*_e_ = 28-31°C (estimated mean = 29.6°C), which revealed the existence of distinct thresholds, or “breakpoints”, in the functional relationship between *T*_th_ and *T*_e_. To the left of each breakpoint (passive warming), the thoracic temperature of the living bees (*T*_th_) was roughly equal to, and varied in unison with, the temperature of the dead bee placed at the same spot and experiencing identical thermal environment (*T*_e_). To the right of breakpoints (active cooling), in contrast, thoracic temperature was lower than, and tended to increasingly diverge from, operative temperature (Fig. 7).

**Fig. 7.**
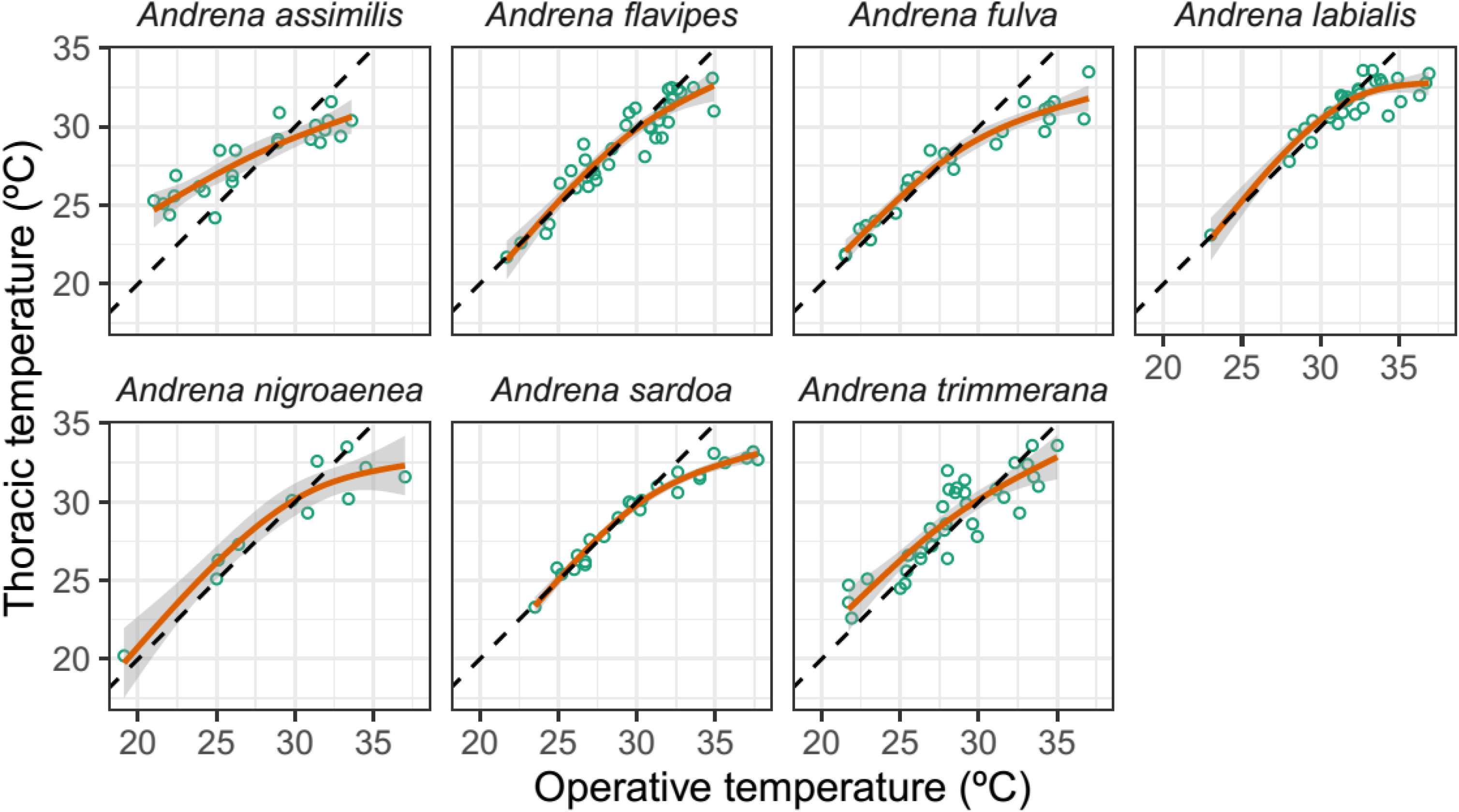
Relationships between thoracic and operative temperatures measured in the field for seven *Andrena* species. Each symbol corresponds to an individual bee, red solid lines are nonparametric regressions fitted using smoothing splines and the shaded bands are 95% confidence envelopes around the fit. The dashed line depicts the *y* = *x* line.

### Comparisons with other bees

Estimates of the warming constant (*K*) for the species of *Andrena* studied here were significantly higher (*F*_1,29_ = 15.01, *P* = 0.0006), and tended to decline more slowly with body mass (*F*_1,29_ = 4.68, *P* = 0.038; interaction term in the linear model), than those of eight bee species from other genera sampled in the study region (Fig. 8A). As a consequence of the different slopes of the declining relationships of *K* with body mass, the divergence between *Andrena* and non-*Andrena* bee species in warming constant tended to broaden as body mass increased (Fig. 8A).

**Fig. 8.**
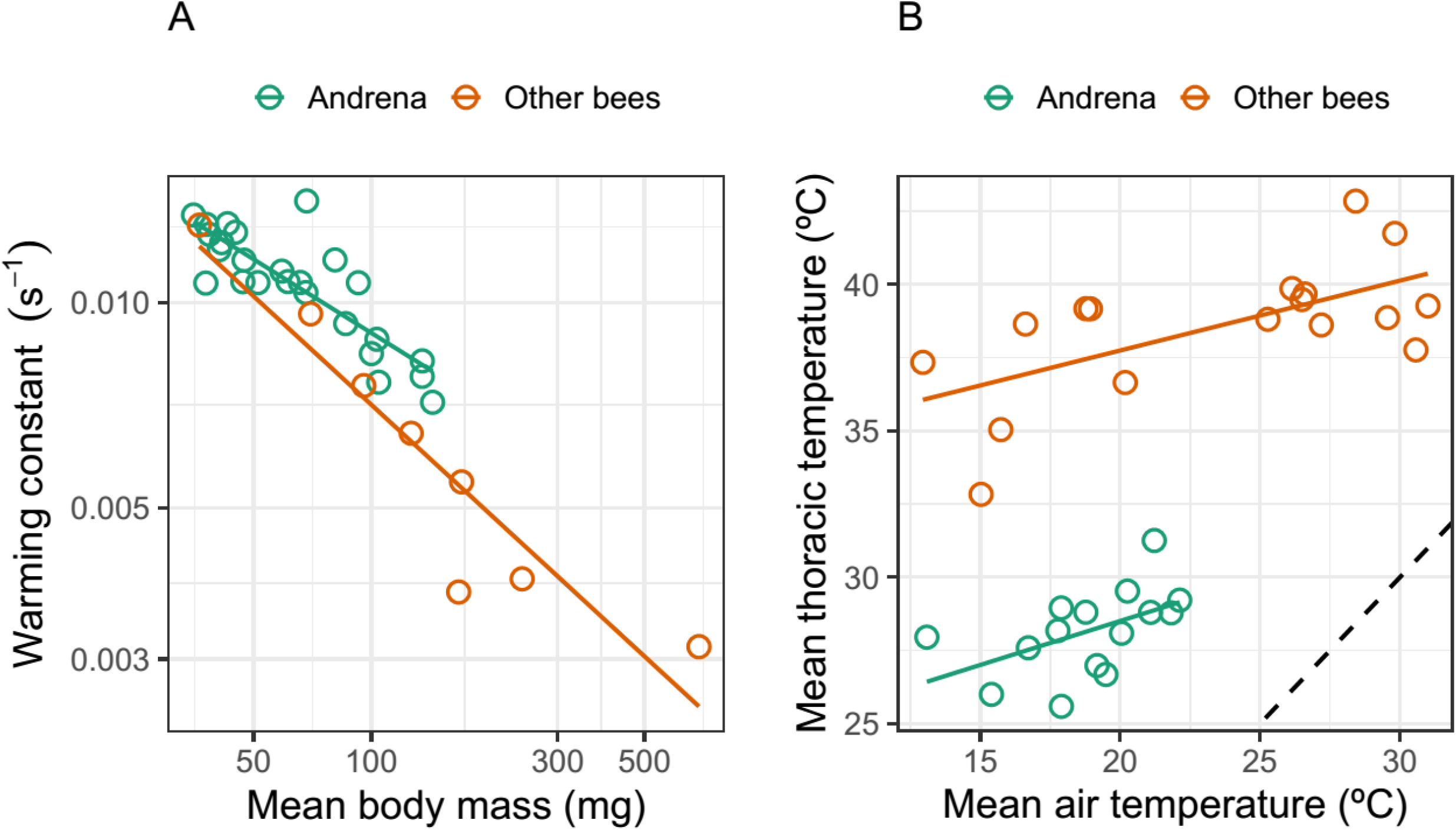
A comparison of the warming constant (A; note logarithmic scale on both axes) and mean thoracic temperature (B) of the *Andrena* species considered in this study with the corresponding values for bees from other genera and families sampled in the same study area and measured using identical laboratory and field methods. Solid lines are fitted linear regressions. The dashed line in B depicts the *y* = *x* line. See Appendix S2: Table S1 for species names, raw data and sources for non-*Andrena* bees included in the graphs.

Mean thoracic temperature measured on foraging bees in the field (*T*_th_) was significantly lower for all species of *Andrena* than for the sample of 17 species of bees from other genera measured in the same region (*F*_1,28_ = 211.14, *P* = 1.4e-14) (Fig. 8B). The relationship across species between mean *T*_th_ and mean air temperature (*T*_a_) was roughly parallel for *Andrena* and non-*Andrena* species, as denoted by the statistical nonsignificance of the interaction term in the linear model (*F*_1,28_ = 0.11, *P* = 0.74). Mean *T*_th_ of *Andrena* species was on average (± SE) 10.5 ± 3.7°C lower than mean *T*_th_ of non-*Andrena* ones over the entire *T*_a_ range represented in the sample (Fig. 8B).

## Discussion

At the plant community level, the numerical importance of *Andrena* bees as pollinators was strongly seasonal, reaching a maximum among those plant species which flowered in late winter and early spring. This pattern was largely independent of seasonal changes in the taxonomic composition of plant species in bloom. *Andrena* pollinators were recorded only during the cooler half, and were absent during the warmer half, of the annual range of air temperatures experienced at flowers by the entire bee assemblage. These ecological restrictions of *Andrena* bees, and the ensuing community-level pattern in plant-pollinator assembly, can be related to distinctive features of their thermal biology: higher warming constants than other bees; null or negligible endothermy; ability to forage at lower body temperatures than bees with endothermic ability; low upper tolerable limit of body temperature beyond which thermal stress probably precluded foraging at the warmest season of the year; and weak thermoregulatory capacity, as shown by the steep linear relationship between thoracic and air temperature and the nearly coincident variation of body and operative temperature over most of the thermal range. These thermal features and their functional relationships, as well as some of their ecological implications in relation to the assembly of plant-pollinator communities and climate change are discussed below.

### Seasonality: cool-blooded bees for early-blooming plants

Results of the present study corroborate for a whole plant community the earlier findings obtained with smaller plant species samples in other habitats of Eurasia and North America, which likewise revealed an association between early blooming plants and pollination by *Andrena* (see references in Introduction). When the whole regional community of entomophilous plants from our Sierra de Cazorla area was considered, *Andrena* bees emerged as numerically important bee pollinators for the subset of species which flowered during the earliest period of the community-wide flowering season. Early-flowering, *Andrena*-pollinated plants were mostly herbs in the families Amaryllidaceae, Brassicaceae, Geraniaceae, Ranunculaceae and Rosaceae (Appendix S1: Table S1) from open, sunny habitats whose flowers had open, non-restrictive corollas with pollen and nectar easily accessible by short-tongued insects. The association between *Andrena* bees and these plants for pollination, however, is unlikely to reflect mutual adaptations evolved in the particular ecological context of our study. On one side, plant flowering times and bee activity periods are both phylogenetically conserved traits at the genus and family levels (Kochmer and Handel 1986, Davies et al. 2013, Turley et al. 2022). And on the other, the early season peak in proportional importance of *Andrena* as pollinators persisted after statistically accounting for effects of plant phylogeny (using families and genera as proxies) on the frequency of *Andrena* as pollinators. It must be stressed here that the generalized linear mixed models fitted to *Andrena* frequency in bee pollinator assemblages of individual plant species did reveal non-zero variance components associated with plant families and genera. This supports a phylogenetic signal of *Andrena* pollination in the plant community studied which would deserve further study.

In accordance with an activity period predominantly restricted to late winter and spring, the thermal range experienced by *Andrena* bees while foraging at flowers (*T*_a_ = ~10-25°C) was substantially narrower, and had a much lower upper limit, than the thermal range experienced by all bees over the whole flowering season (*T*_a_ = ~10-37°C). Since it has been sometimes emphasized that endothermy is a crucial adaptive response of bees to northern or cool climates (Stone and Willmer 1989, Heinrich 1993, Bishop and Armbruster 1999, Gérard et al. 2018), it would have been expected that the association between *Andrena* bees and cool microclimates in the montane habitats studied should be linked to widespread endothermy. Our results falsify this expectation and illustrate that successful exploitation of cooler microclimates by bees does not require endothermy, provided that bees possess the ability to fly at the low body temperatures attainable exclusively by passive, ectothermic warming. In *Andrena*, however, this latter feature apparently comes with a restriction of foraging activity at higher ambient temperatures, which explains their absence from the warmer parts of the seasonal thermal spectrum as discussed in the next section.

### Thermal biology

We characterized the thermal biology of many species of *Andrena* from different subgenera, yet field and laboratory results were remarkably homogeneous across species. In the field, the range and average values of thoracic temperatures for individual species fell within rather narrow limits, and the shape of the relationships between *T*_th_ and *T*_a_ were similar for all species, as were also the critical upper thresholds above which *T*_th_ shifted from roughly equal (passive warming) to lower than operative temperature (active cooling). In the laboratory experiments, differences between species in endothermic ability and warming constant (*K*) were parsimoniously explained by variation in body size, two relationships which have been long known for insects (May 1976, Willmer and Unwin 1981, Stone and Willmer 1989, Bishop and Armbruster 1999). Since at least several of the subgenera included in our species sample most likely are monophyletic lineages (e.g., Chlorandrena, Lepidandrena, Melandrena, Simandrena; Bossert et al. 2022), the prevailing homogeneity of thermal features across our species sample suggests that they represent evolutionarily conserved characteristics within the genus *Andrena* which probably apply to congeneric species from other continents and ecological scenarios as well. This is supported by the low minimum temperatures for flight reported by Stone and Willmer (1989) for three British species, including two of those studied here (*A. clarkella, A. fulva, A. nigroaenea*); the low takeoff thoracic temperatures and insignificant temperature elevation index found by Bishop and Armbruster (1999) for two species from Alaska (*A. thaspii*, *A. frigida*); and the reported inability of central European *Andrena taraxaci* to autonomously raise thoracic temperature under experimental conditions (Schmaranzer et al. 1997). The thermal features of *Andrena* reported here will most likely explain their seasonality in other plant-pollinator systems from the Holarctic realm as well (see references in Introduction).

Mean thoracic temperatures (*T*_th_) were lower, and mean warming constants (*K*) were higher, for *Andrena* than for other bees from 11 different genera sampled in the same area after controlling for variation in body size and air temperature, respectively. The vast majority of non-*Andrena* species used for comparisons belonged to the families Apidae and Megachilidae (Appendix 2: Table S1), all of which are presumably endothermic (Stone and Willmer 1989, Bishop and Armbruster 1999). Therefore, the broad disparity between *Andrena* and non-*Andrena* bees in mean *T*_th_ is just what would be expected from any comparison between ectothermic and endothermic bees (Stone and Willmer 1989, Bishop and Armbruster 1999). The contrast between the two groups in intrinsic heat transfer properties, however, whereby ectothermic *Andrena* had higher *K* values than endothermic species of other genera, was not anticipated and, as far as we know, has been not reported previously. *K* provides an estimate of the rate of heat gain of an object placed in a warmer environment or, alternatively, of the rate of heat loss when placed in a cooler environment. The higher *K* values of *Andrena* thus suggest an intrinsic ability to passively warm or cool significantly faster than other bees for the same body mass, which should have the obvious benefit of reducing thermal inertia and allowing faster adjustment of body temperature when moving across within-habitat mosaics of ambient temperatures. The structural or mechanistic basis of the higher *K* values shown by *Andrena* remains to be investigated, but it seems plausible to suggest that enhanced heat transfer in this genus should be related to some combination of coat color, body geometry, reflectance and hairiness (May 1976, Willmer and Unwin 1981, Pereboom and Biesmeijer 2003).

Other key thermal features of *Andrena* include their inability to significantly warm up autonomously, and the low mean and low upper limit of thoracic temperatures. For the same air temperature, mean thoracic temperature was ~10°C lower for *Andrena* than for endothermic bees (Fig. 8). The ranges of mean thoracic temperatures for *Andrena* (26-31°C) and other bees (33-43°C) did not overlap, and the upper limit for *Andrena* matched the critical thermal threshold beyond which *T*_th_ shifted from passive warming to active cooling (~30-31°C). The *T*_th_ *vs. T*_a_ regression for *Andrena* species (Fig. 8) leads to the projection that, on average, the critical *T*_th_ thermal ceiling of ~30-31°C would be reached at an ambient temperature ~25°C, remarkably close to the temperature above which *Andrena* pollinators virtually vanished from flowers in our study area (Fig. 3). When taken together, these observations strongly support the interpretation that the absence of *Andrena* bees from the latter half of the community-wide flowering season was imposed by their intrinsic inability to withstand thoracic temperatures above ~31°C for long periods without overheating. We tentatively suggest that muscular power output, foraging activity and/or foraging efficiency would be severely depressed above this upper threshold (Coelho 1991, Sinclair et al. 2016), and that lowering thoracic temperature below operative temperature by behavioral means for long periods would severely reduce efficient foraging and constrain microhabitat selection (see Herrera 1995 for supporting data from *Andrena bicolor*).

### *Ectothermy* vs. *endothermy in bees: ecological implications*

Studies on bee thermal biology have historically focused on a few, phylogenetically closely-related lineages of large bees in the family Apidae possessing outstanding endothermic and thermoregulatory ability (e.g., genera *Anthophora, Apis, Bombus, Xylocopa*; see references in Introduction). Generalizations on the thermal biology of bees, including claims on the universality of endothermy, the latter’s importance to colonize northern or cool climates, or its significance for the pollination of early-blooming or cool-climate plants, have been largely grounded on pioneering work on large endothermic bees, particularly bumble bees (Heinrich 1974, 1975, 1993). Two lines of evidence, however, suggest that such generalizations were premature. On one hand, thermal biology seems to have a strong phylogenetic component in bees, with different evolutionary lineages differing in body temperature, tolerable temperature limits and minimum temperature required for flight (Stone and Willmer 1989, Stone 1994, Bishop and Armbruster 1999). Generalizations based on phylogenetically biased species samples are thus prone to bias as well. And on the other hand, the few investigations which have so far examined the thermal biology of a modest number of bee species from three major families other than Apidae (Colletidae, Andrenidae, Halictidae; Stone and Willmer 1989, Herrera 1995, Potts 1995, Bishop and Armbruster 1999, and present study) have consistently failed to support significant endothermy. The combined number of species in these three families (Colletidae, ~2600 species; Andrenidae, ~3000; Halictidae, ~4500; Danforth et al. 2019) amount to a half of the ~20,000 bee species worldwide (Ascher and Pickering 2020) and nearly doubles that of the most-frequently investigated family (Apidae, ~6000 species). It seems plausible to predict, therefore, that future studies on bee thermal biology encompassing a broader, less phylogenetically biased spectrum of species will disclose that bee endothermy is far less frequent than generally assumed to date, and that ectothermic bees comprise a sizeable fraction of bee pollinator communities worldwide.

Energetic constraints have been long acknowledged to drive interactions between plants and bees for pollination (Heinrich 1975, 1981, McCallum et al. 2013). Flower choice and foraging behavior of bees are constrained by the need of balancing foraging costs and the energy returns gained from pollen and nectar (Kadmon & Shmida 1992, Higginson et al. 2006, Willmer 2011). In comparison to ectothermic species, endothermic ones incur additional energetic costs derived from raising body temperature by metabolic means (Heinrich 1974, 1993), which will most likely set the two groups apart with regard to their relationships with plants for pollination (Heinrich 1975, 1981, Herrera 1995, 1997, McCallum et al. 2013). For example, differences in thoracic temperatures between endothermic and ectothermic bees will most likely lead to differences in muscular power output, foraging speed and flower visitation rates (Herrera 1989, 1997). Contrasting thermal biologies (ectothermic cool-blooded *vs*. endothermic hot-blooded) could represent two distinct adaptive pathways allowing bees to exploit distinct segments of the seasonal thermal gradient and associated floral resources. Evaluating this hypothesis will require assessing the proportions of these two thermally distinct groups of bee pollinators in different plant-pollinator systems, and conducting further research on the thermal biology of quantitatively important worldwide but hitherto little-explored bee lineages, as exemplified by the present study on *Andrena* (see also Stone and Willmer 1989, Stone 1994, Herrera 1995, Potts 1995, Bishop and Armbruster 1999).

### Ectothermic bees and climate warming

In addition to helping to understand the mechanisms driving the assembly of plant-pollinator systems along environmental gradients, further research on the thermal biology of under-studied but species-rich families and genera of bees are bound to contribute new insights on the heterogeneity of bee responses to climate warming. Recent reviews and empirical research on the impact of climate change on insect pollinators have often acknowledged the importance of thermal biology as a determinant of bee responses to climate warming, but most inferences have been based on studies of large-sized endothermic species, particularly bumble bees in the genus *Bombus* (Scaven and Rafferty 2013, Ploquin et al 2013, Rasmont et al. 2015, Oyen et al. 2016, Marshall et al 2020, Ghisbain et al. 2021). In the same vein, concerns on wild bee declines have been almost invariably based on data from just two endothermic genera (*Apis*, *Bombus*) which collectively account for a tiny fraction of all species of bees (~1.5%) and whose biological and ecological attributes are far from representative of wild bees in most respects (Wood et al. 2020, Ghisbain 2021). Results of this study suggest that medium- and large-sized ectothermic bees with low upper thermal limits and weak thermoregulatory ability can be more adversely affected by increasing ambient temperatures than the large endothermic species with high thermoregulatory ability most frequently investigated. For example, the increase in the annual number of days with maximum daily temperature >25°C which is underway in our study region as a consequence of climate warming (C. M. Herrera, *personal observations*) will reduce the total annual time available for foraging by *Andrena* bees. In contrast, accelerated climate warming in our study region (Coll et al. 2017, Carvalho et al. 2021, Peña-Angulo et al. 2021) is apparently favoring the increase of small ectothermic bees in pollinator communities (e.g., species of *Heriades, Halictus* subgen. Seladonia, *Lasioglossum* subgen. Evylaeus, *Panurgus*; Herrera 2019: Fig. 10). Attaining a more balanced and realistic apprehension of the impact of climate warming on bees and their pollinating services will require a taxonomically broader and biologically deeper understanding of their thermal biology, particularly of the most species-rich, but nearly uninvestigated ectothermic lineages.

## Acknowledgments

We are indebted to Consejería de Medio Ambiente, Junta de Andalucía, for granting permission to work in the Sierra de Cazorla and providing invaluable facilities there; Klaus Schönitzer and Thomas Wood for assistance with specimen identification; and Pilar Bazaga and Esmeralda López-Perea for technical assistance. Initial funding for this work was provided by grants PB87-0452, PB91-0114 and PB96-0856 from Dirección General de Investigación Científica y Técnica (DGICYT). During its latest stages we were supported by Ministerio de Ciencia e Innovación through European Regional Development Fund (SUMHAL, LIFEWATCH-2019-09-CSIC-13) and Consejería de Transformación Económica, Industria, Conocimiento y Universidades, Junta de Andalucía (P18-FR-4413).

## Author contributions

C. M. H. conceptualized the study, performed field and laboratory work, analyzed the data and led writing. A. N. and L. O. A. identified all bee specimens considered in the study and contributed to the interpretation of results. C. A. provided financial support and project management during the latest stages of the study, and contributed to the analysis of data and interpretation of results. All authors participated in data assembly and curation, edited and revised manuscript drafts and approved the final version.

