## Appendix S1 for "Seasonality of pollinators in Mediterranean montane habitats: cool-blooded bees for early-blooming plants"

**Appendix S1. Plant and bee species sampled.**

TABLE S1. List of the 275 plant species from the Sierra de Cazorla region, southeastern Spain, sampled for pollinator composition, with mean sampling date, total sampling effort, and proportional importance of *Andrena* individuals recorded in pollinator censuses relative to individuals of all bee species combined.

| | | | | Individuals recorded | | |
|---|---|---|---|---|---|---|
| Plant species | Family | Mean census date (days from 1 January) | Number of 3-min pollinator censuses | *Andrena* | All bees | Proportion of *Andrena* |
| *Achillea odorata* | Asteraceae | 175 | 147 | 3 | 23 | 0.130 |
| *Acinos alpinus* | Lamiaceae | 154 | 94 | 0 | 34 | 0.000 |
| *Alliaria petiolata* | Brassicaceae | 127 | 77 | 2 | 3 | 0.667 |
| *Allium guttatum* | Amaryllidaceae | 196 | 90 | 3 | 31 | 0.097 |
| *Allium roseum* | Amaryllidaceae | 140 | 205 | 0 | 26 | 0.000 |
| *Allium rouyi* | Amaryllidaceae | 201 | 115 | 0 | 47 | 0.000 |
| *Allium scorodoprasum* | Amaryllidaceae | 212 | 215 | 3 | 152 | 0.020 |
| *Allium sphaerocephalon* | Amaryllidaceae | 196 | 85 | 8 | 88 | 0.091 |
| *Alyssum montanum* | Brassicaceae | 132 | 105 | 19 | 19 | 1.000 |
| *Alyssum serpyllifolium* | Brassicaceae | 123 | 90 | 0 | 2 | 0.000 |
| *Alyssum simplex* | Brassicaceae | 108 | 85 | 0 | 1 | 0.000 |
| *Amelanchier ovalis* | Rosaceae | 145 | 75 | 8 | 15 | 0.533 |
| *Anagallis monelli* | Primulaceae | 170 | 100 | 12 | 58 | 0.207 |
| *Anarrhinum laxiflorum* | Plantaginaceae | 175 | 50 | 0 | 16 | 0.000 |
| *Andryala integrifolia* | Asteraceae | 163 | 105 | 4 | 73 | 0.055 |
| *Andryala ragusina* | Asteraceae | 201 | 70 | 5 | 58 | 0.086 |
| *Anthemis pedunculata* | Asteraceae | 153 | 95 | 0 | 9 | 0.000 |
| *Anthericum baeticum* | Asparagaceae | 171 | 76 | 11 | 42 | 0.262 |
| *Anthyllis ramburii* | Fabaceae | 167 | 65 | 6 | 8 | 0.750 |
| *Anthyllis vulneraria* | Fabaceae | 144 | 172 | 6 | 77 | 0.078 |
| *Antirrhinum australe* | Plantaginaceae | 172 | 80 | 0 | 7 | 0.000 |
| *Aphyllanthes monspeliensis* | Asparagaceae | 157 | 200 | 1 | 24 | 0.042 |
| *Aquilegia pyrenaica* | Ranunculaceae | 163 | 354 | 1 | 113 | 0.009 |
| *Aquilegia vulgaris* | Ranunculaceae | 151 | 298 | 3 | 132 | 0.023 |

| | | | | | | |
|---|---|---|---|---|---|---|
| *Arabis auriculata* | Brassicaceae | 105 | 74 | 0 | 0 | |
| *Arabis verna* | Brassicaceae | 100 | 101 | 0 | 1 | 0.000 |
| *Arbutus unedo* | Ericaceae | 365 | 110 | 0 | 13 | 0.000 |
| *Arenaria armerina* | Caryophyllaceae | 195 | 96 | 0 | 10 | 0.000 |
| *Arenaria grandiflora* | Caryophyllaceae | 143 | 105 | 3 | 8 | 0.375 |
| *Arenaria modesta* | Caryophyllaceae | 162 | 110 | 0 | 0 | |
| *Arenaria obtusiflora* | Caryophyllaceae | 169 | 100 | 3 | 12 | 0.250 |
| *Armeria filicaulis* | Plumbaginaceae | 153 | 161 | 11 | 21 | 0.524 |
| *Armeria villosa* | Plumbaginaceae | 170 | 75 | 3 | 8 | 0.375 |
| *Asphodelus cerasiferus* | Asphodelaceae | 145 | 136 | 33 | 183 | 0.180 |
| *Astragalus bourgaeanus* | Fabaceae | 133 | 85 | 1 | 7 | 0.143 |
| *Astragalus incanus* | Fabaceae | 131 | 78 | 0 | 37 | 0.000 |
| *Atropa baetica* | Solanaceae | 182 | 70 | 0 | 54 | 0.000 |
| *Bellis perennis* | Asteraceae | 137 | 110 | 4 | 22 | 0.182 |
| *Bellis sylvestris* | Asteraceae | 121 | 100 | 13 | 28 | 0.464 |
| *Berberis hispanica* | Berberidaceae | 159 | 175 | 17 | 46 | 0.370 |
| *Biscutella laxa* | Brassicaceae | 124 | 120 | 5 | 6 | 0.833 |
| *Buxus sempervirens* | Buxaceae | 86 | 130 | 1 | 3 | 0.333 |
| *Calamintha nepeta* | Lamiaceae | 233 | 80 | 0 | 33 | 0.000 |
| *Campanula dieckii* | Campanulaceae | 176 | 130 | 28 | 56 | 0.500 |
| *Campanula mollis* | Campanulaceae | 205 | 110 | 0 | 17 | 0.000 |
| *Campanula rotundifolia* | Campanulaceae | 172 | 87 | 5 | 10 | 0.500 |
| *Carduncellus monspelliensium* | Asteraceae | 180 | 115 | 2 | 50 | 0.040 |
| *Carduus platypus* | Asteraceae | 156 | 65 | 0 | 3 | 0.000 |
| *Carduus tenuiflorus* | Asteraceae | 168 | 81 | 1 | 53 | 0.019 |
| *Carlina hispanica* | Asteraceae | 228 | 145 | 0 | 118 | 0.000 |
| *Carlina racemosa* | Asteraceae | 238 | 140 | 0 | 218 | 0.000 |
| *Carthamus lanatus* | Asteraceae | 197 | 90 | 0 | 38 | 0.000 |
| *Catananche caerulea* | Asteraceae | 172 | 150 | 1 | 118 | 0.008 |
| *Centaurea calcitrapa* | Asteraceae | 189 | 126 | 0 | 204 | 0.000 |
| *Centaurea castellanoides* | Asteraceae | 202 | 135 | 0 | 94 | 0.000 |
| *Centaurea graminifolia* | Asteraceae | 177 | 105 | 2 | 88 | 0.023 |
| *Centaurium tenuiflorum* | Gentianaceae | 175 | 90 | 0 | 1 | 0.000 |
| *Cerastium gibraltaricum* | Caryophyllaceae | 163 | 85 | 1 | 8 | 0.125 |
| *Chaenorhinum macropodum* | Plantaginaceae | 155 | 85 | 0 | 7 | 0.000 |
| *Chiliadenus glutinosus* | Asteraceae | 216 | 83 | 0 | 72 | 0.000 |
| *Chondrilla juncea* | Asteraceae | 229 | 175 | 1 | 239 | 0.004 |
| *Cirsium acaule* | Asteraceae | 225 | 105 | 0 | 23 | 0.000 |
| *Cirsium monspessulanum* | Asteraceae | 222 | 75 | 0 | 68 | 0.000 |
| *Cirsium odontolepis* | Asteraceae | 218 | 100 | 0 | 133 | 0.000 |
| *Cirsium pyrenaicum* | Asteraceae | 224 | 140 | 0 | 70 | 0.000 |

| | | | | | | |
|---|---|---|---|---|---|---|
| *Cirsium vulgare* | Asteraceae | 227 | 110 | 0 | 102 | 0.000 |
| *Cistus albidus* | Cistaceae | 113 | 100 | 22 | 122 | 0.180 |
| *Cistus monspeliensis* | Cistaceae | 152 | 154 | 26 | 100 | 0.260 |
| *Cistus salviifolius* | Cistaceae | 114 | 81 | 36 | 80 | 0.450 |
| *Clematis vitalba* | Ranunculaceae | 197 | 96 | 1 | 27 | 0.037 |
| *Cleonia lusitanica* | Lamiaceae | 168 | 130 | 3 | 86 | 0.035 |
| *Clinopodium vulgare* | Lamiaceae | 191 | 70 | 0 | 24 | 0.000 |
| *Conopodium arvense* | Apiaceae | 160 | 110 | 16 | 17 | 0.941 |
| *Convolvulus arvensis* | Convolvulaceae | 193 | 80 | 0 | 21 | 0.000 |
| *Convolvulus boissieri* | Convolvulaceae | 159 | 100 | 2 | 18 | 0.111 |
| *Coris monspeliensis* | Primulaceae | 166 | 80 | 0 | 25 | 0.000 |
| *Cotoneaster granatensis* | Rosaceae | 159 | 80 | 6 | 8 | 0.750 |
| *Crataegus monogyna* | Rosaceae | 175 | 90 | 11 | 23 | 0.478 |
| *Crepis albida* | Asteraceae | 158 | 67 | 0 | 54 | 0.000 |
| *Crepis capillaris* | Asteraceae | 137 | 95 | 0 | 46 | 0.000 |
| *Crepis vesicaria* | Asteraceae | 161 | 95 | 44 | 90 | 0.489 |
| *Crocus nevadensis* | Iridaceae | 39 | 90 | 23 | 29 | 0.793 |
| *Crocus nudiflorus* | Iridaceae | 274 | 100 | 0 | 36 | 0.000 |
| *Cuscuta triumvirati* | Convolvulaceae | 199 | 105 | 0 | 32 | 0.000 |
| *Cytisus reverchonii* | Fabaceae | 144 | 100 | 5 | 33 | 0.152 |
| *Daphne gnidium* | Thymelaeaceae | 227 | 100 | 0 | 8 | 0.000 |
| *Daphne laureola* | Thymelaeaceae | 93 | 107 | 0 | 10 | 0.000 |
| *Digitalis obscura* | Plantaginaceae | 161 | 180 | 2 | 224 | 0.009 |
| *Draba hispanica* | Brassicaceae | 115 | 102 | 5 | 7 | 0.714 |
| *Drimia maritima* | Hyacinthaceae | 236 | 220 | 1 | 235 | 0.004 |
| *Echinospartum boissieri* | Fabaceae | 174 | 95 | 6 | 31 | 0.194 |
| *Echium flavum* | Boraginaceae | 146 | 160 | 1 | 97 | 0.010 |
| *Erica arborea* | Ericaceae | 99 | 95 | 4 | 22 | 0.182 |
| *Erinacea anthyllis* | Fabaceae | 145 | 136 | 6 | 71 | 0.085 |
| *Erodium cazorlanum* | Geraniaceae | 158 | 75 | 2 | 4 | 0.500 |
| *Erodium cheilanthifolium* | Geraniaceae | 158 | 105 | 16 | 19 | 0.842 |
| *Erodium cicutarium* | Geraniaceae | 115 | 91 | 4 | 8 | 0.500 |
| *Erodium primulaceum* | Geraniaceae | 82 | 100 | 2 | 12 | 0.167 |
| *Erophila verna* | Brassicaceae | 94 | 90 | 5 | 5 | 1.000 |
| *Eryngium campestre* | Apiaceae | 223 | 250 | 0 | 199 | 0.000 |
| *Eryngium dilatatum* | Apiaceae | 198 | 200 | 12 | 180 | 0.067 |
| *Erysimum cazorlense* | Brassicaceae | 148 | 125 | 1 | 1 | 1.000 |
| *Erysimum mediohispanicum* | Brassicaceae | 153 | 100 | 7 | 19 | 0.368 |
| *Erysimum myriophyllum* | Brassicaceae | 140 | 100 | 0 | 9 | 0.000 |
| *Euphorbia nicaeensis* | Euphorbiaceae | 174 | 140 | 10 | 72 | 0.139 |
| *Filipendula vulgaris* | Rosaceae | 171 | 72 | 0 | 3 | 0.000 |
| *Fritillaria lusitanica* | Liliaceae | 107 | 90 | 22 | 35 | 0.629 |
| *Fumana baetica* | Cistaceae | 169 | 160 | 0 | 25 | 0.000 |

| | | | | | | |
|---|---|---|---|---|---|---|
| *Fumana paradoxa* | Cistaceae | 182 | 106 | 3 | 14 | 0.214 |
| *Fumana procumbens* | Cistaceae | 175 | 101 | 1 | 11 | 0.091 |
| *Gagea soleirolii* | Liliaceae | 116 | 108 | 5 | 14 | 0.357 |
| *Galatella linosyris* | Asteraceae | 262 | 120 | 0 | 32 | 0.000 |
| *Galium verum* | Rubiaceae | 194 | 85 | 2 | 5 | 0.400 |
| *Genista longipes* | Fabaceae | 160 | 112 | 9 | 17 | 0.529 |
| *Genista pseudopilosa* | Fabaceae | 172 | 100 | 4 | 14 | 0.286 |
| *Genista scorpius* | Fabaceae | 103 | 112 | 18 | 43 | 0.419 |
| *Geranium cataractarum* | Geraniaceae | 167 | 90 | 1 | 13 | 0.077 |
| *Geranium lucidum* | Geraniaceae | 122 | 104 | 3 | 10 | 0.300 |
| *Geranium molle* | Geraniaceae | 150 | 102 | 8 | 18 | 0.444 |
| *Geum sylvaticum* | Rosaceae | 123 | 122 | 10 | 19 | 0.526 |
| *Gladiolus illyricus* | Iridaceae | 175 | 190 | 3 | 98 | 0.031 |
| *Globularia spinosa* | Plantaginaceae | 141 | 110 | 1 | 5 | 0.200 |
| *Globularia vulgaris* | Plantaginaceae | 101 | 110 | 0 | 10 | 0.000 |
| *Gymnadenia conopsea* | Orchidaceae | 160 | 110 | 0 | 0 | |
| *Halimium atriplicifolium* | Cistaceae | 141 | 40 | 5 | 23 | 0.217 |
| *Hedera helix* | Araliaceae | 276 | 153 | 1 | 63 | 0.016 |
| *Helianthemum apenninum* | Cistaceae | 150 | 165 | 14 | 48 | 0.292 |
| *Helianthemum asperum* | Cistaceae | 147 | 96 | 10 | 18 | 0.556 |
| *Helianthemum cinereum* | Cistaceae | 153 | 100 | 1 | 41 | 0.024 |
| *Helianthemum oelandicum* | Cistaceae | 145 | 167 | 13 | 34 | 0.382 |
| *Helleborus foetidus* | Ranunculaceae | 63 | 1786 | 13 | 200 | 0.065 |
| *Hepatica nobilis* | Ranunculaceae | 109 | 90 | 8 | 10 | 0.800 |
| *Himantoglossum hircinum* | Orchidaceae | 147 | 66 | 1 | 9 | 0.111 |
| *Hippocrepis bourgaei* | Fabaceae | 149 | 95 | 0 | 4 | 0.000 |
| *Hormathophylla baetica* | Brassicaceae | 112 | 41 | 3 | 3 | 1.000 |
| *Hormathophylla spinosa* | Brassicaceae | 176 | 90 | 40 | 48 | 0.833 |
| *Hyacinthoides reverchonii* | Hyacinthaceae | 124 | 105 | 9 | 14 | 0.643 |
| *Hypericum caprifolium* | Hypericaceae | 198 | 76 | 0 | 45 | 0.000 |
| *Hypericum ericoides* | Hypericaceae | 199 | 80 | 0 | 1 | 0.000 |
| *Hypericum perforatum* | Hypericaceae | 193 | 140 | 9 | 91 | 0.099 |
| *Hypochaeris radicata* | Asteraceae | 175 | 60 | 1 | 13 | 0.077 |
| *Iberis carnosa* | Brassicaceae | 106 | 125 | 9 | 9 | 1.000 |
| *Inula montana* | Asteraceae | 170 | 155 | 4 | 198 | 0.020 |
| *Iris foetidissima* | Iridaceae | 170 | 90 | 0 | 27 | 0.000 |
| *Iris planifolia* | Iridaceae | 68 | 100 | 1 | 9 | 0.111 |
| *Jasonia tuberosa* | Asteraceae | 218 | 70 | 0 | 61 | 0.000 |
| *Jonopsidium prolongoi* | Brassicaceae | 121 | 97 | 9 | 10 | 0.900 |
| *Jurinea humilis* | Asteraceae | 166 | 92 | 0 | 15 | 0.000 |
| *Klasea nudicaulis* | Asteraceae | 171 | 75 | 1 | 37 | 0.027 |
| *Klasea pinnatifida* | Asteraceae | 169 | 165 | 0 | 81 | 0.000 |
| *Knautia subscaposa* | Caprifoliaceae | 167 | 177 | 25 | 61 | 0.410 |

| | | | | | | |
|---|---|---|---|---|---|---|
| *Lamium amplexicaule* | Lamiaceae | 112 | 76 | 3 | 30 | 0.100 |
| *Lavandula latifolia* | Lamiaceae | 220 | 818 | 0 | 464 | 0.000 |
| *Leontodon longirrostris* | Asteraceae | 174 | 65 | 0 | 56 | 0.000 |
| *Leopoldia comosa* | Hyacinthaceae | 123 | 100 | 0 | 15 | 0.000 |
| *Lepidium petrophilum* | Brassicaceae | 112 | 115 | 6 | 6 | 1.000 |
| *Leucanthemopsis pallida* | Asteraceae | 132 | 70 | 4 | 31 | 0.129 |
| *Linaria aeruginea* | Plantaginaceae | 155 | 71 | 0 | 4 | 0.000 |
| *Linaria verticillata* | Plantaginaceae | 163 | 75 | 0 | 7 | 0.000 |
| *Linaria viscosa* | Plantaginaceae | 155 | 105 | 0 | 45 | 0.000 |
| *Linum appressum* | Linaceae | 150 | 100 | 0 | 22 | 0.000 |
| *Linum bienne* | Linaceae | 177 | 172 | 76 | 89 | 0.854 |
| *Linum narbonense* | Linaceae | 158 | 85 | 0 | 10 | 0.000 |
| *Linum tenue* | Linaceae | 171 | 190 | 6 | 109 | 0.055 |
| *Lithodora fruticosa* | Boraginaceae | 154 | 185 | 37 | 79 | 0.468 |
| *Lonicera arborea* | Caprifoliaceae | 173 | 65 | 1 | 63 | 0.016 |
| *Lotus corniculatus* | Fabaceae | 164 | 90 | 0 | 44 | 0.000 |
| *Lysimachia ephemerum* | Primulaceae | 183 | 200 | 0 | 38 | 0.000 |
| *Lythrum baeticum* | Lythraceae | 172 | 85 | 0 | 49 | 0.000 |
| *Lythrum junceum* | Lythraceae | 166 | 97 | 0 | 52 | 0.000 |
| *Lythrum salicaria* | Lythraceae | 238 | 110 | 0 | 66 | 0.000 |
| *Mantisalca salmantica* | Asteraceae | 199 | 160 | 0 | 79 | 0.000 |
| *Marrubium supinum* | Lamiaceae | 164 | 160 | 2 | 125 | 0.016 |
| *Mentha aquatica* | Lamiaceae | 228 | 110 | 0 | 42 | 0.000 |
| *Merendera montana* | Colchicaceae | 248 | 120 | 0 | 38 | 0.000 |
| *Muscari neglectum* | Hyacinthaceae | 105 | 93 | 1 | 19 | 0.053 |
| *Narcissus bujei* | Amaryllidaceae | 65 | 205 | 39 | 83 | 0.470 |
| *Narcissus cuatrecasasii* | Amaryllidaceae | 113 | 214 | 10 | 45 | 0.222 |
| *Narcissus hedraeanthus* | Amaryllidaceae | 86 | 205 | 58 | 86 | 0.674 |
| *Narcissus longispathus* | Amaryllidaceae | 90 | 295 | 55 | 206 | 0.267 |
| *Narcissus triandrus* | Amaryllidaceae | 113 | 125 | 4 | 15 | 0.267 |
| *Nepeta tuberosa* | Lamiaceae | 172 | 60 | 0 | 40 | 0.000 |
| *Omphalodes linifolia* | Boraginaceae | 146 | 100 | 1 | 42 | 0.024 |
| *Ononis pusilla* | Fabaceae | 158 | 79 | 8 | 29 | 0.276 |
| *Ononis spinosa* | Fabaceae | 204 | 100 | 0 | 40 | 0.000 |
| *Onopordum acaulon* | Asteraceae | 165 | 115 | 0 | 37 | 0.000 |
| *Onosma tricerosperma* | Boraginaceae | 178 | 90 | 0 | 37 | 0.000 |
| *Orchis coriophora* | Orchidaceae | 175 | 120 | 1 | 37 | 0.027 |
| *Origanum virens* | Lamiaceae | 198 | 77 | 0 | 30 | 0.000 |
| *Ornithogalum umbellatum* | Hyacinthaceae | 142 | 160 | 41 | 91 | 0.451 |
| *Orobanche haenseleri* | Orobanchaceae | 180 | 96 | 0 | 46 | 0.000 |
| *Papaver dubium* | Papaveraceae | 177 | 85 | 6 | 64 | 0.094 |
| *Parentucellia latifolia* | Orobanchaceae | 134 | 33 | 0 | 0 | |
| *Parnassia palustris* | Celastraceae | 263 | 70 | 0 | 0 | |

| | | | | | | |
|---|---|---|---|---|---|---|
| *Petrorhagia nanteuilii* | Caryophyllaceae | 178 | 70 | 0 | 21 | 0.000 |
| *Phlomis herba-venti* | Lamiaceae | 183 | 150 | 0 | 84 | 0.000 |
| *Phlomis lychnitis* | Lamiaceae | 178 | 151 | 0 | 93 | 0.000 |
| *Picnomon acarna* | Asteraceae | 233 | 125 | 0 | 138 | 0.000 |
| *Pilosella pseudopilosella* | Asteraceae | 176 | 66 | 1 | 68 | 0.015 |
| *Pistorinia hispanica* | Crassulaceae | 170 | 161 | 0 | 8 | 0.000 |
| *Plumbago europaea* | Plumbaginaceae | 226 | 205 | 0 | 146 | 0.000 |
| *Polygala boissieri* | Polygalaceae | 146 | 100 | 0 | 6 | 0.000 |
| *Polygonatum odoratum* | Asparagaceae | 142 | 138 | 0 | 11 | 0.000 |
| *Potentilla caulescens* | Rosaceae | 225 | 125 | 0 | 39 | 0.000 |
| *Potentilla reptans* | Rosaceae | 132 | 100 | 7 | 24 | 0.292 |
| *Primula acaulis* | Primulaceae | 111 | 153 | 2 | 18 | 0.111 |
| *Prolongoa hispanica* | Asteraceae | 165 | 88 | 0 | 19 | 0.000 |
| *Prunella laciniata* | Lamiaceae | 174 | 115 | 1 | 19 | 0.053 |
| *Prunus mahaleb* | Rosaceae | 137 | 120 | 11 | 39 | 0.282 |
| *Prunus prostrata* | Rosaceae | 127 | 90 | 0 | 20 | 0.000 |
| *Pterocephalus spathulatus* | Caprifoliaceae | 193 | 120 | 0 | 6 | 0.000 |
| *Ptilostemon hispanicus* | Asteraceae | 224 | 65 | 0 | 74 | 0.000 |
| *Ranunculus bulbosus* | Ranunculaceae | 155 | 100 | 9 | 32 | 0.281 |
| *Ranunculus ficaria* | Ranunculaceae | 127 | 108 | 1 | 3 | 0.333 |
| *Ranunculus gramineus* | Ranunculaceae | 122 | 110 | 4 | 17 | 0.235 |
| *Ranunculus malessanus* | Ranunculaceae | 120 | 95 | 8 | 25 | 0.320 |
| *Ranunculus paludosus* | Ranunculaceae | 131 | 105 | 18 | 38 | 0.474 |
| *Ranunculus repens* | Ranunculaceae | 167 | 100 | 3 | 73 | 0.041 |
| *Rhagadiolus edulis* | Asteraceae | 165 | 95 | 9 | 32 | 0.281 |
| *Roemeria argemone* | Papaveraceae | 145 | 90 | 0 | 29 | 0.000 |
| *Rosa canina* | Rosaceae | 160 | 75 | 40 | 84 | 0.476 |
| *Rosa micrantha* | Rosaceae | 181 | 145 | 16 | 106 | 0.151 |
| *Rosa sicula* | Rosaceae | 175 | 80 | 13 | 45 | 0.289 |
| *Rosmarinus officinalis* | Lamiaceae | 140 | 166 | 7 | 159 | 0.044 |
| *Rubus ulmifolius* | Rosaceae | 191 | 95 | 6 | 191 | 0.031 |
| *Salvia lavandulifolia* | Lamiaceae | 185 | 72 | 0 | 51 | 0.000 |
| *Salvia verbenaca* | Lamiaceae | 133 | 90 | 0 | 39 | 0.000 |
| *Santolina rosmarinifolia* | Asteraceae | 192 | 130 | 2 | 29 | 0.069 |
| *Saponaria ocymoides* | Caryophyllaceae | 153 | 141 | 0 | 0 | |
| *Satureja intricata* | Lamiaceae | 193 | 147 | 0 | 56 | 0.000 |
| *Saxifraga carpetana* | Saxifragaceae | 134 | 90 | 1 | 16 | 0.063 |
| *Saxifraga haenseleri* | Saxifragaceae | 132 | 77 | 2 | 2 | 1.000 |
| *Saxifraga tridactylites* | Saxifragaceae | 123 | 60 | 0 | 0 | |
| *Scabiosa andryaefolia* | Caprifoliaceae | 182 | 62 | 0 | 10 | 0.000 |
| *Scilla paui* | Hyacinthaceae | 121 | 90 | 2 | 14 | 0.143 |
| *Scorzonera albicans* | Asteraceae | 160 | 70 | 0 | 1 | 0.000 |
| *Scorzonera cf_reverchonii* | Asteraceae | 150 | 80 | 5 | 9 | 0.556 |

| | | | | | | |
|---|---|---|---|---|---|---|
| *Scorzonera laciniata* | Asteraceae | 133 | 100 | 22 | 22 | 1.000 |
| *Sedum acre* | Crassulaceae | 169 | 100 | 4 | 9 | 0.444 |
| *Sedum album* | Crassulaceae | 185 | 165 | 2 | 65 | 0.031 |
| *Sedum dasyphyllum* | Crassulaceae | 193 | 110 | 0 | 21 | 0.000 |
| *Sedum mucizonia* | Crassulaceae | 172 | 100 | 0 | 24 | 0.000 |
| *Sedum sediforme* | Crassulaceae | 200 | 95 | 0 | 77 | 0.000 |
| *Senecio doria* | Asteraceae | 222 | 74 | 0 | 40 | 0.000 |
| *Senecio malacitanus* | Asteraceae | 299 | 115 | 7 | 50 | 0.140 |
| *Seseli montanum* | Apiaceae | 266 | 120 | 0 | 15 | 0.000 |
| *Sherardia arvensis* | Rubiaceae | 117 | 80 | 0 | 0 | |
| *Sideritis incana* | Lamiaceae | 178 | 130 | 0 | 73 | 0.000 |
| *Silene colorata* | Caryophyllaceae | 155 | 92 | 0 | 4 | 0.000 |
| *Silene psammitis* | Caryophyllaceae | 149 | 132 | 0 | 5 | 0.000 |
| *Sisymbrella aspera* | Brassicaceae | 171 | 180 | 6 | 42 | 0.143 |
| *Sisymbrium crassifolium* | Brassicaceae | 146 | 70 | 6 | 12 | 0.500 |
| *Solidago virgaurea* | Asteraceae | 235 | 100 | 0 | 19 | 0.000 |
| *Sonchus aquatilis* | Asteraceae | 232 | 91 | 0 | 106 | 0.000 |
| *Stachys officinalis* | Lamiaceae | 190 | 138 | 0 | 70 | 0.000 |
| *Taraxacum laevigatum* | Asteraceae | 145 | 120 | 11 | 45 | 0.244 |
| *Taraxacum obovatum* | Asteraceae | 96 | 122 | 3 | 47 | 0.064 |
| *Teucrium aureum* | Lamiaceae | 178 | 161 | 6 | 58 | 0.103 |
| *Teucrium rotundifolium* | Lamiaceae | 176 | 181 | 0 | 42 | 0.000 |
| *Teucrium webbianum* | Lamiaceae | 192 | 81 | 0 | 35 | 0.000 |
| *Thymus mastichina* | Lamiaceae | 169 | 160 | 10 | 93 | 0.108 |
| *Thymus orospedanus* | Lamiaceae | 153 | 163 | 2 | 29 | 0.069 |
| *Thymus serpylloides* | Lamiaceae | 168 | 75 | 0 | 37 | 0.000 |
| *Trifolium campestre* | Fabaceae | 179 | 82 | 26 | 31 | 0.839 |
| *Ulex parviflorus* | Fabaceae | 88 | 125 | 2 | 83 | 0.024 |
| *Valeriana tuberosa* | Caprifoliaceae | 136 | 165 | 17 | 39 | 0.436 |
| *Verbascum giganteum* | Scrophulariaceae | 218 | 115 | 0 | 44 | 0.000 |
| *Verbascum sinuatum* | Scrophulariaceae | 217 | 115 | 0 | 41 | 0.000 |
| *Verbena officinalis* | Verbenaceae | 183 | 120 | 0 | 111 | 0.000 |
| *Viburnum tinus* | Adoxaceae | 111 | 90 | 4 | 14 | 0.286 |
| *Vicia onobrychioides* | Fabaceae | 162 | 65 | 0 | 54 | 0.000 |
| *Vicia pseudocracca* | Fabaceae | 154 | 73 | 0 | 35 | 0.000 |
| *Vicia sativa* | Fabaceae | 140 | 100 | 0 | 12 | 0.000 |
| *Viola cazorlensis* | Violaceae | 149 | 153 | 0 | 0 | |
| *Viola odorata* | Violaceae | 108 | 242 | 0 | 35 | 0.000 |

TABLE S2. List of the 58 species of bees for which air temperature was measured at foraging sites and number of measurements taken per species.

| Bee species | Number of measurements |
|---|---|
| *Amegilla albigena* | 27 |
| *Amegilla ochroleuca* | 21 |
| *Amegilla quadrifasciata* | 15 |
| *Andrena* unidentified | 4 |
| *Andrena assimilis* | 23 |
| *Andrena baetica* | 27 |
| *Andrena bicolor* | 71 |
| *Andrena bimaculata* | 2 |
| *Andrena flavipes* | 37 |
| *Andrena fulva* | 24 |
| *Andrena humilis* | 1 |
| *Andrena labialis* | 32 |
| *Andrena livens* | 26 |
| *Andrena nigroaenea* | 11 |
| *Andrena pilipes* | 1 |
| *Andrena rhenana* | 2 |
| *Andrena sardoa* | 26 |
| *Andrena schencki* | 1 |
| *Andrena thoracica* | 1 |
| *Andrena tibialis* | 3 |
| *Andrena trimmerana* | 35 |
| *Andrena vulpecula* | 1 |
| *Anthidiellum breviusculum* | 51 |
| *Anthidium florentinum* | 82 |
| *Anthophora aestivalis* | 3 |
| *Anthophora affinis* | 1 |
| *Anthophora crassipes* | 33 |
| *Anthophora dispar* | 64 |
| *Anthophora fulvodimidiata* | 1 |
| *Anthophora leucophaea* | 3 |
| *Anthophora plumipes* | 1 |

| | |
|---|---|
| *Anthophora retusa* | 3 |
| *Anthophora romandii* | 14 |
| *Apis mellifera* | 220 |
| *Bombus pascuorum* | 36 |
| *Bombus pratorum* | 30 |
| *Bombus terrestris* | 118 |
| *Bombus vestalis* | 2 |
| *Ceratina chalybea* | 1 |
| *Ceratina cucurbitina* | 3 |
| *Ceratina mocsaryi* | 39 |
| *Colletes cunicularius* | 29 |
| *Eucera elongatula* | 25 |
| *Halictus scabiosae* | 7 |
| *Lasioglossum* unidentified | 1 |
| *Lithurgus chrysurus* | 59 |
| *Megachile albisecta* | 39 |
| *Megachile centuncularis* | 2 |
| *Megachile octosignata* | 1 |
| *Megachile pilidens* | 10 |
| *Melecta italica* | 1 |
| *Nomada succincta* | 1 |
| *Osmia bicornis* | 9 |
| *Osmia cornuta* | 17 |
| *Rhodanthidium sticticum* | 3 |
| *Xylocopa cantabrita* | 3 |
| *Xylocopa valga* | 5 |
| *Xylocopa violacea* | 19 |
