## Appendix S2 for "Seasonality of pollinators in Mediterranean montane habitats: cool-blooded bees for early-blooming plants"

**Appendix S2. Thermal traits of non-*Andrena* bees.**

TABLE S1. Non-*Andrena* bee species, body mass and thermal data used in the analyses presented in the section *Results: Comparisons with other bees*, and plotted in Fig. 8.

| Species | Family | Mean body mass (mg) | Warming constant, *K* ($s^{-1}$) | Mean thoracic temperature (ºC) | Mean air temperature (ºC) | Source |
|---|---|---|---|---|---|---|
| *Amegilla albigena* (Lepeletier, 1841) | Apidae | | | 39.3 | 31.0 | Herrera (1997) |
| *Amegilla ochroleuca* (Pérez, 1879) | Apidae | | | 39.7 | 26.6 | Herrera (1997) |
| *Amegilla quadrifasciata* (Curtis, 1829) | Apidae | | | 41.8 | 29.8 | Herrera (1997) |
| *Anthidium florentinum* (Fabricius, 1775) | Megachilidae | 245.4 | | 38.9 | 29.6 | Herrera (1997) |
| *Anthophora crassipes* Lepeletier, 1841 | Apidae | | | 38.6 | 27.2 | Herrera (1997) |
| *Anthophora dispar* Lepeletier, 1841 | Apidae | 167.2 | 0.00376 | 38.7 | 16.6 | C. M. Herrera (*personal observation*) |
| *Anthophora plumipes* (Pallas, 1772) | Apidae | 130.1 | | 35.0 | 15.7 | C. M. Herrera (*personal observation*) |

| | | | | | | |
|---|---|---|---|---|---|---|
| *Anthophora romandii* Lepeletier, 1841 | Apidae | 93.5 | | 39.2 | 18.8 | C. M. Herrera (*personal observation*) |
| *Apis mellifera* Linnaeus, 1758 | Apidae | 95.6 | 0.00756 | 38.8 | 25.3 | Herrera (1997), C. M. Herrera (*personal observation*) |
| *Bombus pascuorum* (Scopoli, 1763) | Apidae | 170.0 | 0.00546 | 39.5 | 26.5 | Herrera (1997), C. M. Herrera (*personal observation*) |
| *Bombus pratorum* (Linnaeus, 1761) | Apidae | 335.9 | | 37.3 | 12.9 | C. M. Herrera (*personal observation*) |
| *Bombus terrestris* (Linnaeus, 1758) | Apidae | 243.0 | 0.00394 | 39.9 | 26.1 | Herrera (1997), C. M. Herrera (*personal observation*) |
| *Colletes cunicularius* (Linnaeus, 1761) | Colletidae | 126.4 | 0.00643 | | | C. M. Herrera (*personal observation*) |
| *Dasypoda morotei* Quilis, 1928 | Melittidae | 69.9 | 0.00963 | | | C. M. Herrera (*personal observation*) |
| *Megachile pilidens* Alfken, 1924 | Megachilidae | | | 37.8 | 30.6 | Herrera (1997) |
| *Osmia bicornis* (Linnaeus, 1758) | Megachilidae | 57.9 | | 36.7 | 20.2 | C. M. Herrera (*personal observation*) |
| *Osmia cornuta* (Latreille, 1805) | Megachilidae | 89.8 | | 32.8 | 15.0 | C. M. Herrera (*personal observation*) |
| *Panurgus banksianus* (Kirby, 1802) | Andrenidae | 36.3 | 0.01299 | | | C. M. Herrera (*personal observation*) |
| *Xylocopa cantabrita* Lepeletier, 1841 | Apidae | 276.6 | | 39.2 | 18.9 | C. M. Herrera (*personal observation*) |
| *Xylocopa violacea* (Linnaeus, 1758) | Apidae | 688.5 | 0.00313 | 42.9 | 28.4 | Herrera (1997), C. M. Herrera (*personal observation*) |
